# Atlas55+: Brain Functional Atlas of Resting-state Networks for Late Adulthood

**DOI:** 10.1101/2020.07.13.200824

**Authors:** Gaelle E. Doucet, Loic Labache, Paul M. Thompson, Marc Joliot, Sophia Frangou, the Alzheimer’s Disease Neuroimaging Initiative

**Author notes:** Corresponding Author: Dr. Gaelle Doucet, Brain Architecture, Imaging and Cognition Lab, Boys Town National Research Hospital, 555 N 30th street, Omaha, NE 68131, USA. Data used in preparation of this article were obtained from the Alzheimer’s Disease Neuroimaging Initiative (ADNI) database (adni.loni.usc.edu). As such, the investigators within the ADNI contributed to the design and implementation of ADNI and/or provided data but did not participate in analysis or writing of this report. A complete listing of ADNI investigators can be found at: http://adni.loni.usc.edu/wp-content/uploads/how_to_apply/ADNI_Acknowledgement_List.pdf.

## Abstract

Currently, several human brain functional atlases are used to define the spatial constituents of the resting-state networks (RSNs). However, the only brain atlases available are derived from samples of young adults. As brain networks are continuously reconfigured throughout life, the lack of brain atlases derived from older populations may influence RSN results in late adulthood. To address this gap, the aim of the study was to construct a reliable brain atlas derived only from older participants. We leveraged resting-state functional MRI data from three cohorts of healthy older adults (total N=563; age=55-95years) and a younger-adult cohort (N=128; age=18-35 years). We identified the major RSNs and their subdivisions across all older-adult cohorts. We demonstrated high spatial reproducibility of these RSNs with an average spatial overlap of 67%. Importantly, the RSNs derived from the older-adult cohorts were spatially different from those derived from the younger-adult cohort (p=2.3×10^−3^). Lastly, we constructed a novel brain atlas, called Atlas55+, which includes the consensus of the major RSNs and their subdivisions across the older-adult cohorts. Thus, Atlas55+ provides a reliable age-appropriate template for RSNs in late adulthood and is publicly available. Our results confirm the need for age-appropriate functional atlases for studies investigating aging-related brain mechanisms.

## Introduction

Resting-state functional magnetic resonance imaging (rs-fMRI) has been instrumental in mapping the brain functional connectome in health (Fox MD et al. 2005; Smith SM et al. 2009; Doucet G et al. 2011; Smith SM et al. 2015) and disease (Feltz CJ and GE Miller 1996; Buckner RL et al. 2009; Tracy JI and GE Doucet 2015; Doucet GE et al. 2017; Dong D et al. 2018; Sha Z et al. 2019). The brain functional connectome can be defined as a multi-scale partition in which regions may be aggregated into networks, which in turn may be combined into larger systems (Kiviniemi V et al. 2009; Abou Elseoud A et al.2011; Doucet G *et al*. 2011). Previous work by us and others suggest that the intermediate partition of the brain at the level of networks may be optimal to explain cognitive activity and to identify reliable biomarkers of neuropsychiatric and neurological disorders (van den Heuvel MP and HE Hulshoff Pol 2010; Abou Elseoud A *et al*. 2011; Doucet G *et al*. 2011; Doucet G et al. 2012; Doucet GE *et al*. 2017; Salman MS et al. 2019), while limiting multiple comparisons when compared to finer brain partitions (Zalesky A et al. 2010). At this level of brain organization, five major resting-state networks (RSNs) and their subdivisions have been typically identified across rs-fMRI studies: those involved in internally guided, higher order mental functions as part of the intrinsic system (default-mode [DMN], executive control [ECN], and salience [SAL] networks) and those supporting externally driven, specialized sensory and motor processing as part of the extrinsic system (visual [VIS] and sensorimotor [SMN] networks) (De Luca M et al. 2006; Smith SM *et al*. 2009; Doucet G *et al*. 2011; Buckner RL et al. 2013; Doucet GE et al. 2019). Each of these brain networks relies on established white matter pathways (Toosy AT et al. 2004; Greicius MD et al. 2009; van den Heuvel MP et al. 2009) that support the consistency of their spatiotemporal configuration and functional roles (Damoiseaux JS et al. 2006; Smith SM *et al*. 2009; van den Heuvel MP and HE Hulshoff Pol 2010; Doucet G *et al*. 2011; Doucet GE *et al*. 2019; Elliott ML et al. 2019).

Several brain functional atlases are currently available which support reproducibility by harmonizing the RSN definition across neuroimaging studies (Doucet GE *et al*. 2019). However, to date, the only brain functional atlases available are derived from samples of young healthy adults, typically below the age of 40 years (Doucet GE *et al*. 2019). Nonetheless, changes in brain function over the course of adulthood have been well-documented and suggest that brain networks are continuously reconfigured throughout adult life (Damoiseaux JS et al. 2008; Meunier D et al. 2009; He X et al. 2013; Betzel RF et al. 2014; Damoiseaux JS 2017; Varangis E, Q Razlighi, et al. 2019; Yaple ZA et al. 2019; Luo N, J Sui, A Abrol, D Lin, et al. 2020). To our knowledge, there is currently no reference brain functional atlas derived from rs-fMRI data from older adults, and this may undermine neuroimaging efforts to characterize the brain functional connectome and its cognitive role in late adulthood. Critically, differences in the spatial composition of the RSNs derived from large samples of younger versus older adults have yet to be systematically investigated.

In this context, the current study focused on the spatial definition of the five major RSNs (DMN, ECN, SAL, VIS and SMN) and their subdivisions in healthy adults aged 55 years and older. The specific aims were: (a) to test the reproducibility of RSNs in three large independent cohorts comprising a total of 563 older healthy adults (age range: 55-95 years); accordingly, we analyzed data from the Center for Ageing and Neuroscience (CamCAN) project (Shafto MA et al. 2014; Taylor JR et al. 2017), the Southwest University Adult Lifespan Dataset (SALD) (Wei D et al. 2018) and the Alzheimer’s Disease Neuroimaging Initiative (ADNI) database (Jack CR, Jr. et al. 2008; Petersen RC et al. 2010); (b) to identify differences in the anatomical constitution of RSNs defined in the above cohorts compared to those in younger adults (age range 18-35); and (c) generate a reliable consensual RSN atlas to enhance the reproducibility of template-defined RSNs in older adults.

## Materials and Methods

### Cohorts

#### Older-Adult Cohorts

We used data from the CamCAN (Shafto MA *et al*. 2014; Taylor JR *et al*. 2017), the SALD (Wei D *et al*. 2018) and the ADNI datasets (Jack CR, Jr. *et al*. 2008; Petersen RC *et al*. 2010). In each cohort, we selected healthy individuals aged 55 years and above for whom both rs-fMRI and structural MRI data were available. This selection resulted in a sample of 250 individuals for CamCAN (referred to as “CamCAN55+”), 190 for SALD and 134 for ADNI (details in supplementary material). Following quality control of the imaging data (described below), 11 participants across cohorts were removed for excessive head motion. The total analysis sample for the older individuals comprised 563 individuals, described in Table 1 and Supplementary Figure S1.

**Table 1:**
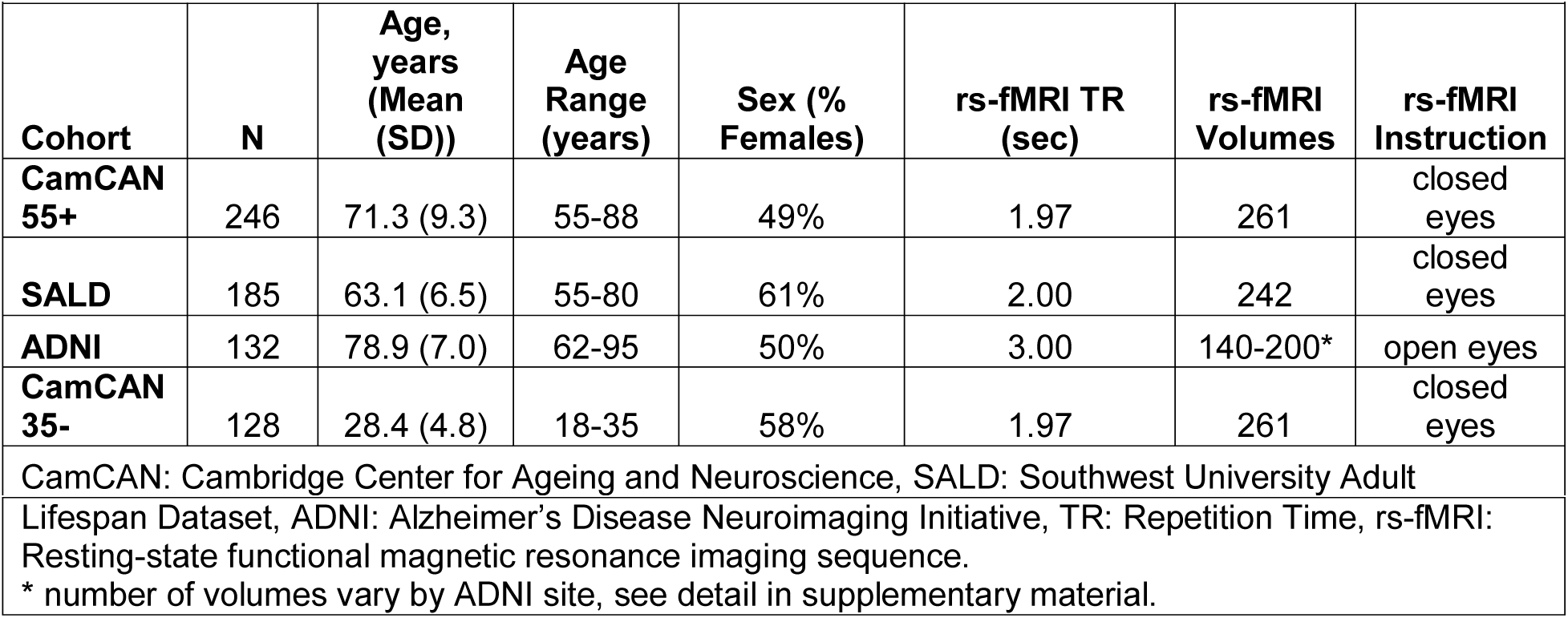
Cohort Characteristics

| Table 1: Cohort Characteristics |  |  |  |  |  |  |  |
| --- | --- | --- | --- | --- | --- | --- | --- |
| Cohort | N | Age, years (Mean (SD)) | Age Range (years) | Sex (% Females) | rs-fMRI TR (sec) | rs-fMRI Volumes | rs-fMRI Instruction |
| CamCAN 55+ | 246 | 71.3 (9.3) | 55-88 | 49% | 1.97 | 261 | closed eyes |
| SALD | 185 | 63.1 (6.5) | 55-80 | 61% | 2.00 | 242 | closed eyes |
| ADNI | 132 | 78.9 (7.0) | 62-95 | 50% | 3.00 | 140-200* | open eyes |
| CamCAN 35- | 128 | 28.4 (4.8) | 18-35 | 58% | 1.97 | 261 | closed eyes |
| CamCAN: Cambridge Center for Ageing and Neuroscience, SALD: Southwest University Adult |  |  |  |  |  |  |  |

|  |
| --- |
| Lifespan Dataset, ADNI: Alzheimer's Disease Neuroimaging Initiative, TR: Repetition Time, rs-fMRI: Resting-state functional magnetic resonance imaging sequence.<br>* number of volumes vary by ADNI site, see detail in supplementary material. |

#### Younger-Adult Cohort

A subset of the CamCAN cohort, referred to as “CamCAN35-”, was used to test for differences between younger (18 to 35 years-old) and older (55 years and older) adults in RSN spatial composition. This age range was chosen to match the typical age range of healthy participants used in currently available functional atlases (Doucet GE *et al*. 2019). The CamCAN35-included 128 individuals. Following the same quality control process, we excluded two individuals from CamCan35- (Table 1).

### Resting-state fMRI Acquisition and Preprocessing

In the CamCAN cohort, rs-fMRI data were acquired while participants rest with their eyes closed on a 3T Siemens TIM Trio scanner with a 32-channel head coil with the following acquisition parameters: TR/TE= 1970/30 msec, 32 axial slices, flip angle =78 degrees; FOV =192 mm×192 mm; voxel-size =3 mm×3 mm×4.44 mm, acquisition time=8min 40sec, number of volumes: 261. More information on the MRI sequences can be found in Taylor JR *et al*. (2017).

In the SALD cohort, rs-fMRI data were acquired while participants rest with their eyes closed on a 3T Siemens Trio scanner, using the following acquisition parameters (Wei D *et al*. 2018): TR/TE= 2000/30 msec, 32 axial slices, flip angle=90 degrees; FOV=220 mm×220 mm; voxel-size =3.4 mm×3.4 mm×4 mm, acquisition time=8min, number of volumes: 242. More information on the MRI sequences can be found in Wei D *et al*. (2018).

In the ADNI cohort, rs-fMRI data were acquired while participants kept their eyes open. Acquisition tool place in 43 sites using 3T scanners [General Electric (GE), Philips or Siemens] and variable acquisition protocols (details in http://adni.loni.usc.edu/methods/documents/) <u>that commonly used</u> a TR=3000 sec, with 140 to 200 volumes. When multiple resting-state fMRI datasets were available for a participant, the first one was selected for analyses.

Regardless of the cohort, the rs-fMRI data were preprocessed in an identical fashion, using SPM12 and the DPABI Toolbox (Yan CG et al. 2016). Preprocessing procedures for the rs-fMRI datasets included removal of the first 3 volumes, motion correction to the first volume with rigid-body alignment; co-registration between the functional scans and the anatomical T1-weighted scan; spatial normalization of the functional images into Montreal Neurological Institute (MNI) stereotaxic standard space; spatial smoothing within the functional mask with a 6-mm at full-width at half-maximum Gaussian kernel; wavelet despiking (Patel AX et al. 2014); linear detrending; and regression of motion parameters and their derivatives (24-parameter model) (Friston KJ et al. 1996), as well as white matter (WM), CSF time series. The WM and CSF signals were computed using a component based noise reduction method (CompCor, 5 principal components) (Behzadi Y et al. 2007). Lastly, bandpass filtering was applied at [0.01-0.1] Hz (Cordes D et al. 2001).

As previously mentioned, across the three older-adult cohorts, we excluded 11 individuals who had excessive head movement based on maximum transient (volume-to-volume) head motion above 2 mm translation or 1 degree rotation. After removing these individuals, we ensured that head motion, based on the mean framewise displacement (Power JD et al. 2012) did not correlate with age (r=0.08). Following the same process, we excluded 2 individuals from the CamCan35-cohort.

### Resting-state Networks in the older-adult cohorts

#### Resting-state networks identification in the older-adult cohorts

We used a process validated by Naveau M et al. (2012) to identify reliable brain networks. First, for each individual within each cohort, single-subject independent component analyses (ICAs) were conducted 20 times with random initialization using the Multivariate Exploratory Linear Optimized Decomposition into Independent Components (MELODIC) software, version 3.15, included in the FMRIB Software Library (FSL) v6.0.3 (http://www.fmrib.ox.ac.uk/fsl) (Smith SM et al. 2004). The number of independent components (ICs) was estimated by Laplace approximation (Minka T 2000). A symmetric approach of the FastICA algorithm (Hyvarinen A 1999) was used to compute the ICAs. Second, for each repetition, we used the multi-scale clustering of individual component algorithm (MICCA) to classify ICs into *N* groups (Naveau M *et al*. 2012). The number of groups was automatically estimated by the algorithm (Naveau M *et al*. 2012). Third, the *Icasso* algorithm (Himberg J et al. 2004) was used to select groups of reproducible ICs (i.e., groups with ICs that were present in at least 50% of the 20 repetitions), which we identified as the “group-level components” (Salman MS *et al*. 2019). Fourth, for each group-level component, a voxel-wise *t*-score map of individual ICs was computed and thresholded using a mixture model (*p*>0.95, (Beckmann CF and SM Smith 2004)). Lastly, we discarded group-level components if their spatial map: (i) mainly covered non-gray matter (i.e., CSF and WM), or (ii) included regions with strong signal attenuation due to susceptibility artefacts (i.e., lower frontal and lower temporal regions) (Supplementary Figures S2-S4).

#### Spatial Overlap of group-level components in older-adult cohorts

Within each cohort, we mapped the spatial overlap across the group-level components by computing an index that quantified the number of group-level components overlapping at each voxel. This voxel-wise index ranged from 1 (no spatial overlap, the voxel is assigned to only one component) to *n* (where *n* is the maximum number of components overlapping at a voxel).

As the goal of the current study was to create a reliable functional atlas of RSNs, we created non-overlapping group-level components, on which we based the remaining analyses. Accordingly, each brain voxel was assigned to the group-level components with the highest *t*-score.

#### Reliability analyses of the group-level components in older-adult cohorts

As the spatial configuration of a brain network is strongly related with its functional connectivity (Bijsterbosch JD et al. 2018), we conducted supplementary analyses to assess the functional reliability of each group-level component, in each cohort, following the methods described in Labache L et al. (2020) (Supplementary Figure S5). In each cohort, we generated a dendrogram based on the average connectivity matrix between the group-level components. The functional reliability of a group-level component depended on the number of times it maintained the same position in the dendrogram of each individual participant of the same cohort. We created a reliability index (Q) for each group-level component, where a score close to zero indicated perfect functional reliability and a high score indicated high functional unreliability (details in supplementary information and supplementary Figure S5). We conducted Tukey’s fences tests to identify and discard these highly unreliable group-level components.

#### Construction of the major RSNs in older-adult cohorts

We chose to focus on the major RSNs defined as: the DMN, the ECN, the SAL, the SMN, and the VIS (Smith SM *et al*. 2009; Doucet G *et al*. 2011; Doucet GE *et al*. 2019) and their subdivisions and examined their spatial definition across the three cohorts.

Following the identification of non-overlapping and functionally reliable group-level components in each cohort, we proceeded to assigning them to the major RSNs of interest, namely the DMN, ECN, SAL, VIS and SMN. As reference for each network, we used their spatial definition in the Consensual Atlas of REsting-state Networks (CAREN) (Doucet GE *et al*. 2019). This atlas was generated by our group and presents the spatial configuration of the major RSNs based on their common properties across the most widely used brain functional atlases and as such it represents the most reproducible spatial features of the major RSNs currently available. This choice was also based on the fact that, to the best of our knowledge, there are currently no validated and reproducible RSNs from rs-fMRI data of a large sample of older healthy individuals that we could use as normative templates.

In each cohort, we computed the Dice coefficient (Dice LR 1945) to quantify the degree of spatial overlap between each group-level component and each of the five aforementioned CAREN RSNs. Components were then assigned to the RSN with which they had the largest overlap (Supplementary Table S1). We applied this process to each cohort separately to create spatial maps of the major RSNs. We were thus able to construct a spatial map for each RSN. We used the Dice coefficient to test the pairwise similarity between the same-labeled RSNs across cohorts. Lastly, to ensure that the results did not depend on the template used (i.e., CAREN), we also used the atlas created by Yeo BT et al. (2011) for the RSN assignment.

### Spatial comparison of the major RSNs between young and older adults

Prior to testing for differences in the spatial constitution of RSNs between older and younger adults we constructed the major RSNs in the CamCAN35-by applying the same procedure as for the older cohorts (Supplementary Figure S12).

We then compared their respective RSNs to those derived from the other two independent cohorts (ADNI and SALD). For this, we used Dice’s coefficient to quantify the spatial similarity of the RSNs in CamCAN35-, and in CamCAN55+ respectively, to those of SALD and ADNI, and compared these pairwise coefficients (i.e., we compared the coefficients resulting from the RSN comparison CamCAN55+ vs. SALD and ADNI to those resulting from the mRSN comparison CamCAN35- vs. SALD and ADNI). This choice of approach was based on two major reasons: (1) among the three cohorts, CamCAN was the largest and also provided more reliable data (having the largest number of rs-fMRI volumes (Elliott ML *et al*. 2019), Table 1); and (2) this approach enabled us to account for differences in analytic approach, site, and MRI acquisition parameters.

### Generation of Atlas55+

The major RSNs identified in each of the three older-adult cohorts were used as input data to create a consensual atlas, named Atlas55+. For this, we used the function *consensus_similarity*.*m* from the Network Community Toolbox (http://commdetect.weebly.com), which constructs a representative partition.

To test the spatial reproducibility of the major RSNs in Atlas55+, we conducted supplementary analyses: (1) we assigned each brain voxel to a network only if at least two of the three older-adult cohorts assigned this voxel to the same network. We then computed a spatial correlation between the two versions of Atlas55+ using Pearson’s correlation analyses. (2) We created a voxel-wise confidence map to quantify the consistency of the network assignment between the RSNs of the atlas and those from each of the three older-adult cohorts. This voxel-wise index ranged from 0 (poor confidence, the network assignment of a voxel in Atlas55+ differed from its network assignment in all three cohorts) to 100% (high confidence, the network assignment of a voxel in Atlas55+ was identical to its network assignment in all three cohorts). Lastly, we compared the RSNs of the atlas to those of CamCAN35-in order to quantify the spatial differences between RSNs from younger adults and older adults, computing Dice coefficients.

To identify the anatomical definition of each major RSN in the atlas, we generated subject-specific versions using a dual regression approach in FSL (Beckmann CF et al. 2009; Nickerson LD et al. 2017). We then entered each network into a one-sample *t* test, adding age, head motion (mean framewise displacement), estimated total intracranial volume (TIV, measured using FreeSurfer v6.0 (http://surfer.nmr.mgh.harvard.edu/)), and site as covariates of no interest. Significant regions were reported at a *p*<0.05 after applying a family wise error (FWE) correction at the voxel level (*T*>5.3, cluster size>20 voxels).

### Subdivisions of the RSNs within Atlas55+

We then focused on the subdivisions of each RSN within Atlas55+ to increase the spatial resolution of the atlas across the three older-adult cohorts.

For each major RSN, we quantified the degree of spatial overlap between the group-level components assigned to that RSN across the three cohorts, using Dice coefficients (Supplementary Table S1). Based on these coefficients, we constructed a dendrogram and identified clusters with high spatial overlap across independent group-level components, with one component from each of the three cohorts (Supplementary Figure S11). Lastly, we created a spatial map for each of these clusters, which displays the spatial characteristics of a subdivision of that RSN (Supplementary Table S3). A voxel was identified as part of that RSN subdivision if it was found in at least two out of the three components. We chose to focus on clusters with contributions from independent components for all three older-adult cohorts because this suggests that these components were reliably found in each cohort and therefore were minimally influenced by acquisition parameters.

### Effects of sex and age on the major five RSNs in the Atlas55+

The effect of age on the spatial distribution of each major RSN in the atlas (extracted from the dual regression) across all individuals (total n=563) was tested by conducting an analysis of covariance, adding covariates of no interest: head motion, TIV, and site. Sex differences in the spatial distribution of the RSNs in the atlas were also assessed using age, head motion, TIV, and site as covariates of no interest. Significant clusters are reported at a *p*<0.05 with FWE correction at the voxel level (*T*>5.3, cluster size>20 voxels).

## Results

### Identification of group-level components in the older-adult cohorts

In the CamCAN55+, SALD and ADNI cohorts, we respectively identified a total of 27, 24 and 23 reproducible group-level components (Supplementary Figures S2-S4 and S8). For the CamCAN55+ cohort, we found that 27% of the voxels were assigned to a single group-level component, while less than 0.5% of the voxels were assigned to a maximum of nine group-level components. The degree of spatial overlap between components was similar for the other two cohorts (Supplementary Figure S7). Regardless of cohort, the regions with the highest overlap were located in the medial posterior cortex, the dorso-medial prefrontal cortex and the lateral parietal cortex (Supplementary Figure S7).

Regardless of cohort, the functional reliability tests showed that all group-level components had a similar degree of stability, with no outlier, across individuals and in each cohort (Supplementary Figure S6). The reliability Q scores of each component did not differ across cohorts (*p*>0.1).

### Construction and Reliability of the major RSNs in the older-adult cohorts

The five RSNs constructed in each older-adult cohort (Figure 1) showed high spatial reproducibility across the three cohorts with an average spatial overlap of 67% (mean range: 66-69%), with the SAL having the lowest score (46%) and the VIS having the highest score (83%) (Figure 2A). Supplementary Figure S9 shows the voxelwise confidence values with respect to the probability of a voxel being assigned to the same-labeled RSN across the three cohorts. The majority of voxels (83.5%) had confidence values over 66%, indicating that at least two out of the three cohorts had the same RSN assignment.

**Figure 1:**
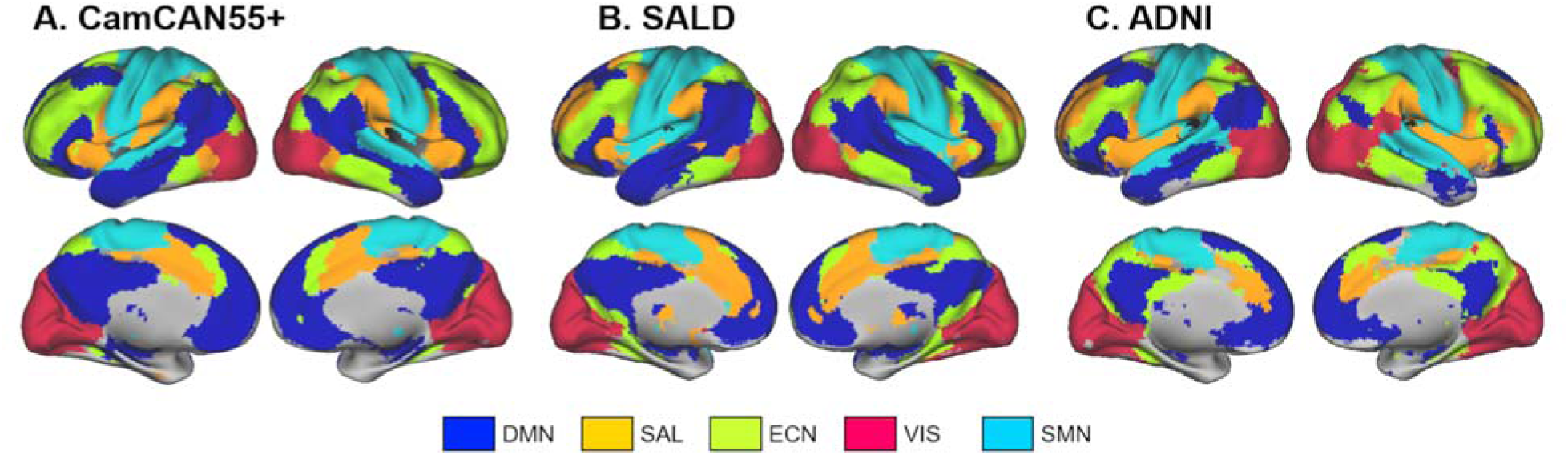
Spatial map of the five major RSNs in each older-adult cohort. (A) CamCAN55+, (B) SALD, (C) ADNI. DMN: Default mode network, SAL: Salience network, ECN: Executive control network, VIS: Visual network, SMN: Sensorimotor network.

**Figure 2:**
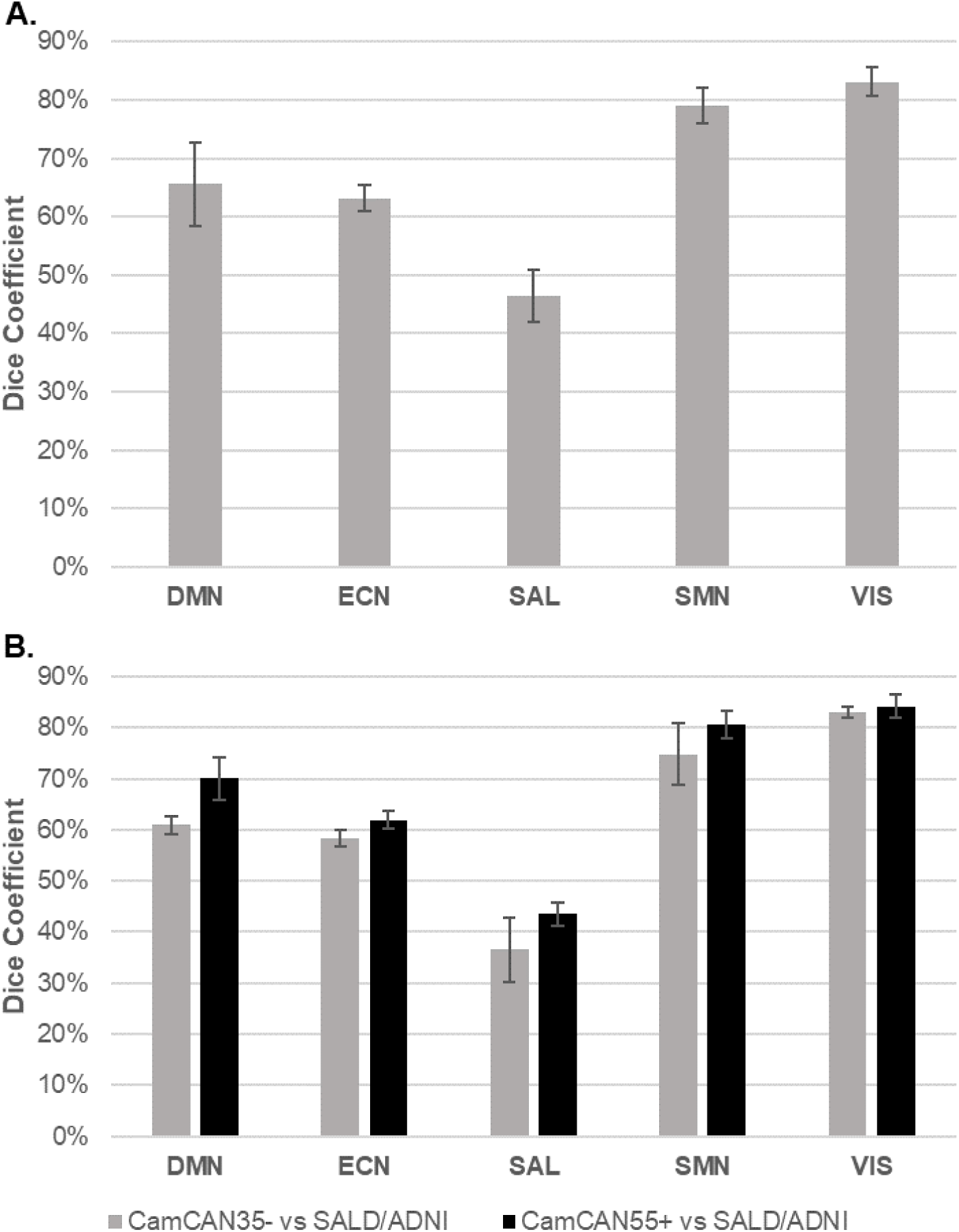
Spatial similarity of the RSNs across the cohorts. (A) Spatial similarity of the RSNs across the older-adult cohorts. (B) Comparison of the spatial similarity of the RSNs identified in the CamCAN35- vs. the ADNI and SALD cohorts and in the CamCAN55+ vs. the ADNI and SALD cohorts. DMN: Default mode network, SAL: Salience network, ECN: Executive control network, VIS: Visual network, SMN: Sensorimotor network.

The choice of a reference atlas, CAREN or Yeo, minimally affected the spatial definition of the RSNs. The average spatial overlap of each RSN constructed based on CAREN versus each RSN constructed based on Yeo’s atlas was 86% across the three cohorts (CamCAN: 86% (sd=15%), SALD: 85% (19%), ADNI: 86% (9%)) (Supplementary Figure S10).

### Comparison of the major RSNs between older- and young adults

We found that the major RSNs in the CamCAN35- (Supplementary Figure S12) were less similar to those identified in the SALD and ADNI than in the CamCAN55+ (t=-4.2, p=2.3×10^−3^; Figure 2B). The SMN and VIS, which cover primary cortices differed the least, while the SAL network showed the largest spatial difference between the younger and older cohorts.

### Atlas55+

Atlas55+ is available at two resolution levels: at the lever of the five major RSNs (DMN, ECN, SAL, SMN and VIS, Figure 3A) and at the level of their 15 subdivisions that were also reliably identified across the three older-adult cohorts (Figure 4). The anatomical description of each RSN and their subdivisions is detailed in Supplementary Tables S2 and S3. The degree of confidence in network assignment in Atlas55+ was high across the three older-adult cohorts (Figure 3B). The SMN and VIS were the most reproducible networks with 77% and 93% of their voxels respectively being consistently assigned across all three cohorts.

**Figure 3:**
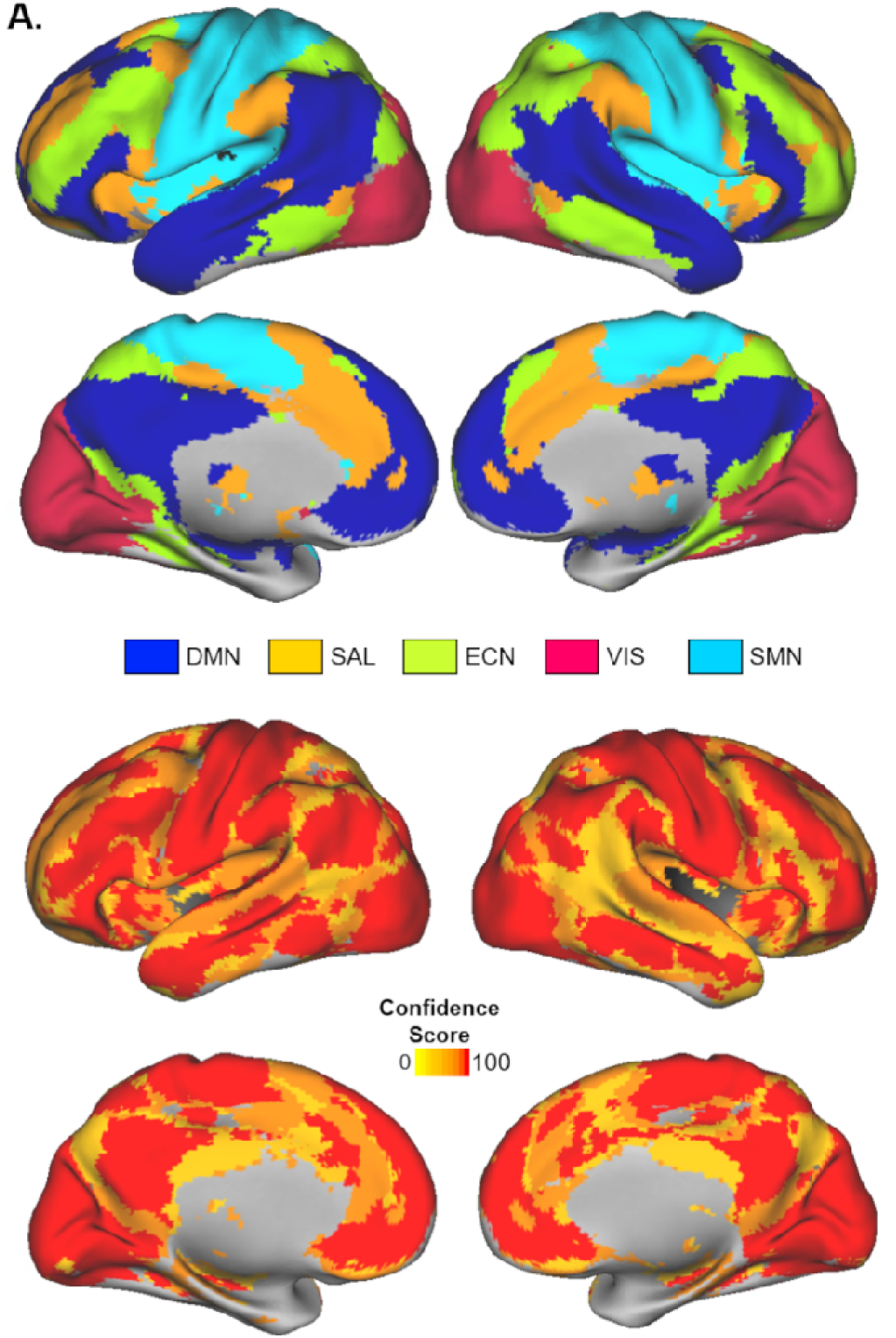
Atlas55+: Consensus brain atlas composed of the five major resting-state networks based on three cohorts of healthy individuals aged 55 years and above. (A) Spatial map of the RSNs. (B) Voxel-wise confidence map in Atlas55+. This measure quantifies the probability that a voxel in an Atlas55+ network is assigned to the same-label RSN in each of the three cohorts. DMN: Default mode network, SAL: Salience network, ECN: Executive control network, VIS: Visual network, SMN: Sensorimotor network.

**Figure 4:**
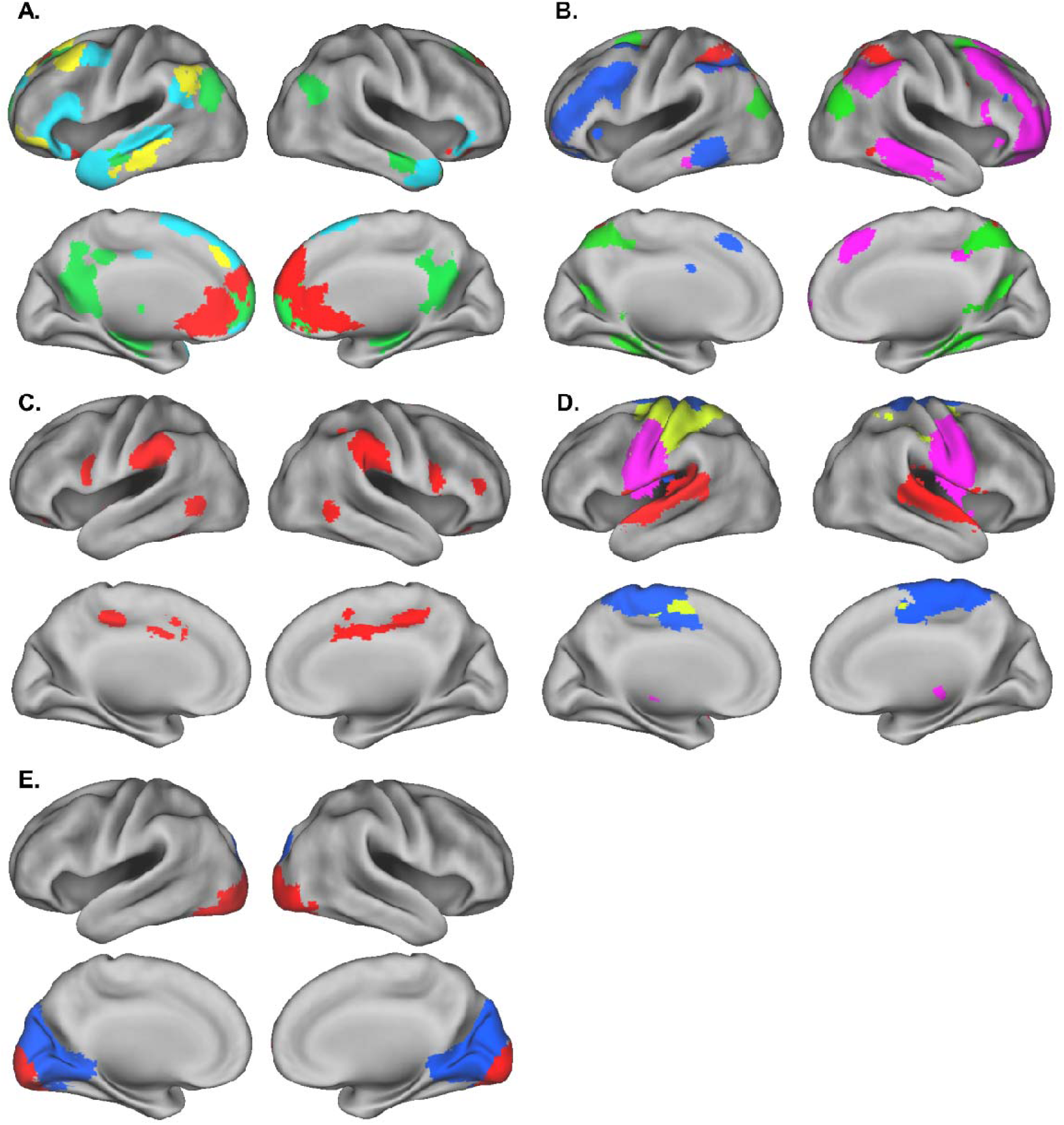
Reliable subdivisions of each of the five major RSNs across the three older-adult cohorts. (A) 4 subdivisions of the DMN; (B) 4 subdivisions of the ECN; (C) 1 subdivision of the SAL; (D) 4 subdivisions of the SMN; (E) 2 subdivisions of the VIS. Each color corresponds to one subdivision. Each subdivision is further described in the supplementary material (Supplementary Table S3 and Supplementary Figure S11).

As defined in Atlas55+, the SMN was comprised of the sensory and motor regions (precentral and postcentral gyrus and supplementary motor area), the primary auditory cortex (superior temporal cortex), and thalamus. The SMN was further partitioned into four subdivisions: the ventral and dorsal parts of the pre- and post-central gyri, the supplementary motor area and the bilateral superior temporal gyri (auditory network). The VIS network largely covered the occipital lobe, and was partitioned into two subdivisions: the medial and posterior parts (Figure 4). Both SMN and VIS networks showed the largest spatial overlap with their respective networks in younger adults as outlined in CamCAN35- (spatial overlap: 69% and 84% respectively) (Figure 5).

**Figure 5:**
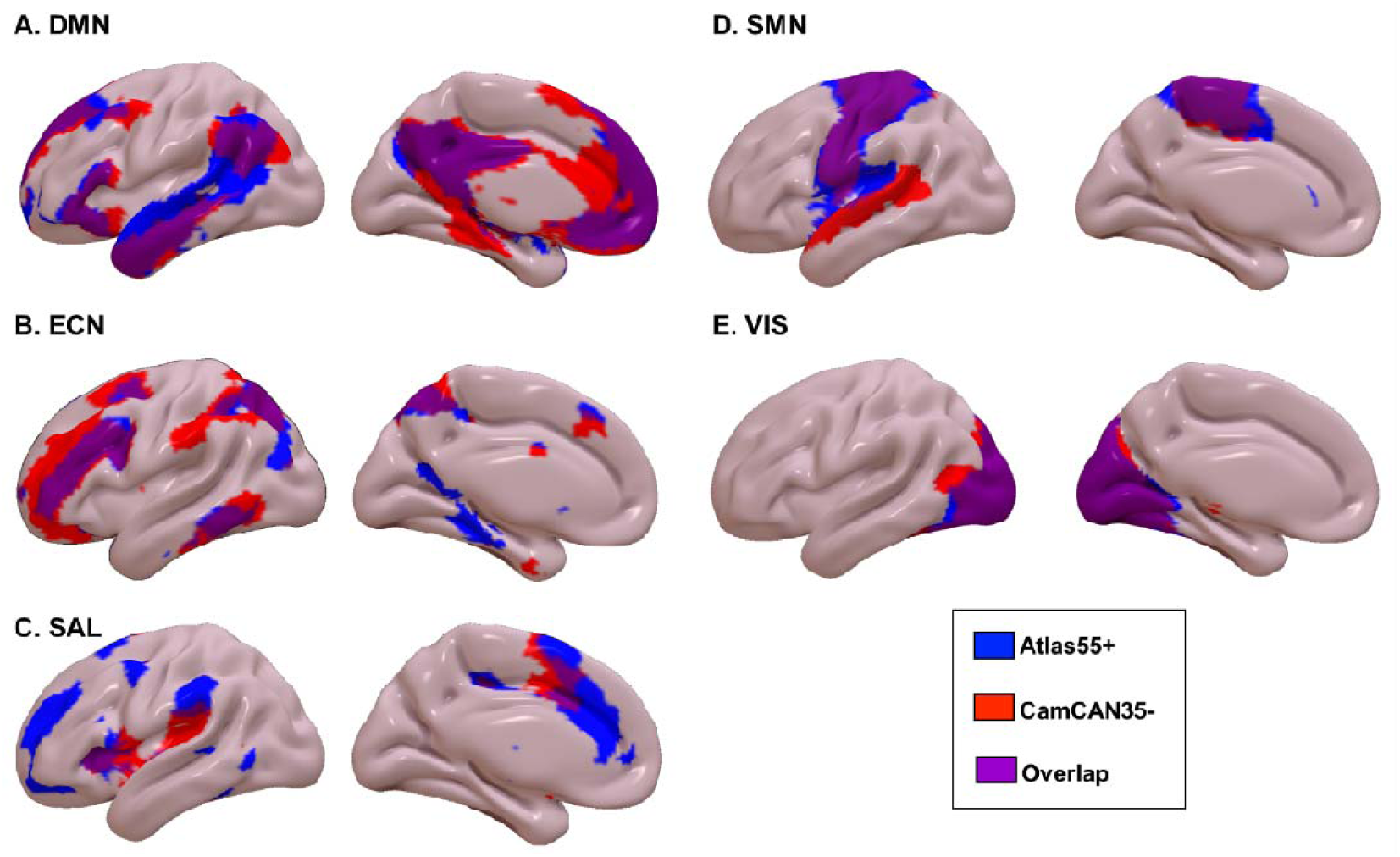
Spatial comparison of the major resting state networks between adults over the age of 55 years, as defined in Atlas55+, and younger adults (aged 18-35 years) from the CamCAN35-cohort.

For the DMN, 51% of the network’s voxels showed consistency across all three older-adult cohorts. In Atlas55+, the DMN comprised the medial prefrontal cortex/ventral anterior cingulate cortex (ACC), the precuneus/posterior cingulate cortex (PCC), the inferior frontal cortex, the angular gyri, the middle temporal cortex, the hippocampi and amygdala, and some bilateral cerebellar clusters. The DMN further included four reliable subdivisions comprising the anterior DMN (ACC), the core DMN (precuneus/PCC, medial prefrontal cortex, bilateral angular gyri), and two left lateralized networks mostly covering the inferior frontal gyrus, temporal poles and the posterior part of the superior temporal gyrus, and dorsal medial prefrontal cortex (Figure 4). The whole DMN showed a 63% overlap with the DMN in younger adults as defined in CamCAN35- (Figure 5). The major spatial differences were localized in the posterior medial temporal lobe -which was part of the DMN from CamCAN35-, but not from Atlas55+ (Figure 5).

For the ECN, 58% of the network’s voxels showed consistency across all three older-adult cohorts. In Atlas55+, the ECN included parts of the dorsolateral prefrontal cortex, the lateral and medial parietal cortex, the posterior inferior temporal cortex, the posterior parahippocampal gyrus and some bilateral cerebellar clusters. The ECN further included four reliable subnetworks comprising the right and left lateral parietal-frontal cortex, as well as two posterior subdivisions mostly covering the posterior parietal lobe and the medial temporal lobe. The whole ECN showed a 60% overlap with the ECN in younger adults as defined in CamCAN35-. The major differences involved the additions of the posterior medial temporal lobe (lingual and parahippocampal gyri) in the ECN from Atlas55+ (Figure 5).

The SAL network emerged as the least reproducible network: 25% of the network’s voxels were consistently assigned to the SAL across the three older-adult cohorts. In Atlas55+, the SAL was comprised of the anterior insula bilaterally, the dorsal ACC, the supramarginal gyri, subcortical regions including the putamen and thalamic nuclei, and bilateral cerebellar clusters. We only found one reliable subdivision which mostly covered the dorsal ACC and supramarginal gyri, bilaterally (Figure 4). The spatial overlap with the whole SAL in younger adults as defined in CamCAN35-was the lowest with 30% and the regional differences were widespread (Figure 5).

### Associations of sex and age with the spatial distribution of the major RSNs in Atlas55+

#### Sex

We did not find any significant differences between males and females on the spatial distribution of any major RSN in the atlas.

#### Age

We found a negative regional association with age for each of the major RSNs in the atlas (Figure 6, Supplementary Table S4). In detail, increasing age was associated with lower spatial integrity of: (i) both anterior and posterior medial regions in the DMN (Figure 6A), (ii) bilateral fronto-parietal regions in the ECN (Figure 6B), (iii) bilateral anterior insula and dorsal anterior cingulate cortex in the SAL (Figure 6C), (iv) bilateral superior temporal and postcentral gyri in the SMN (Figure 6D), and to a lesser degree (v) in the calcarine and lingual gyri of the VIS (Figure 6E).

**Figure 6:**
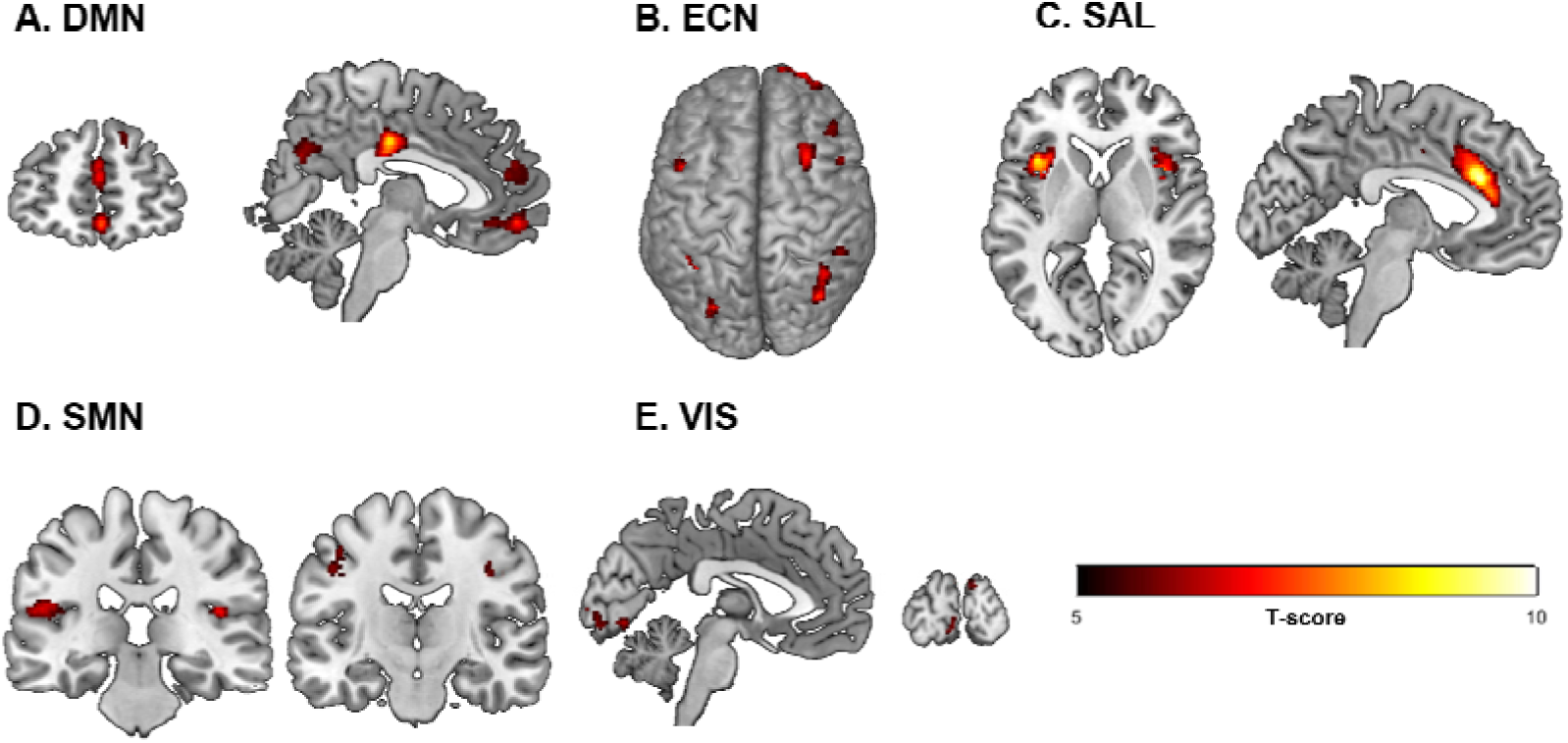
Clusters showing lower spatial integration with the rest of the RSN, with older age, across the three older-adult cohorts. Details of the clusters are in Supplementary Table S3. Significance set up at *p*<0.05 (FWE correction at the voxel level).

In contrast, increasing age was associated with higher spatial integrity of cerebellar regions in both the DMN and the ECN (Supplementary Table S4). No other results were significant.

## Discussion

This study investigated (a) the spatial definition of the major RSNs in healthy individuals in late adulthood using rs-fMRI data from three large independent cohorts of participants aged 55 years and above and; (b) differences in the spatial definition of RSNs in late adulthood to those derived from healthy individuals in mid adulthood. We found a high reproducibility of the spatial definition of the RSNs across the cohorts of older adults, but also significant differences when compared to the RSNs of younger adults. In response to these findings, we constructed Atlas55+, a robust brain functional atlas derived from rs-fMRI of 563 healthy adults between the ages of 55 and 95 years, in order to promote RSN reproducibility in future studies focusing on older populations. The age cut-off of 55 years may be considered arbitrary but there is currently no clear consensus on a specific age that distinguishes middle to late adulthood. In response, we followed the same age criterion as in ADNI, which is a landmark initiative in North America to investigate brain organization in older populations (Jack CR, Jr. *et al*. 2008; Petersen RC *et al*. 2010).

Despite different MRI scanner types, acquisition parameters, and sites, we demonstrated good reproducibility of the spatial definition of the RSNs across the three older-adult cohorts. Regardless of the cohort, the regions with the highest spatial overlap between RSNs were localized in the medial prefrontal cortex, the precuneus, and the lateral parietal cortex, which have been described as brain hubs in young adults (Tomasi & Volkow, 2011; Zuo et al. 2012). Participation in multiple networks is thought to reflect an inherent feature of these associative regions (Yeo BT et al. 2014). While studies have reported a negative effect of aging on the strength of these regional hubs (Damoiseaux JS 2017; Zhang H et al. 2017), the current study suggests that the variability in their location remains relatively unaltered within older healthy populations.

Across the three older-adult cohorts, we found that the most reproducible networks were the SMN and VIS networks. These networks are mostly covering primary cortices, which are known to have high structural-functional coherence (Luo N, J Sui, A Abrol, J Chen, et al. 2020), low inter-individual variability in anatomical morphology (White LE et al. 1997) and in resting-state functional connectivity (Franco AR et al. 2013; Mueller S et al. 2013; Li R et al. 2017), and tend to preferentially participate in single networks in line with their circumscribed and specific functions (Yeo BT *et al*. 2014). In contrast, the SAL network showed the largest spatial differences across the three older-adult cohorts. This was consistent with the fact that we only identified one reliable subdivision of the SAL, in contrast to the other high-order RSNs (i.e., DMN and ECN) that showed four reliable subdivisions each. In studies of individuals below the age of 40 years, the spatial definition and role of the SAL network have been largely variable (Dosenbach NU et al. 2007; Seeley WW et al. 2007; Smith SM *et al*. 2009; Yeo BT *et al*. 2011; Shirer WR et al. 2012; Doucet GE *et al*. 2019) and this appears to be the case in older adults too.

In line with what we hypothesized, we found age-related alterations in the spatial composition of all five RSNs. The association of RSN spatial definition with age was examined in two different and complementary ways. First, by comparing RSNs between older and younger adults and second by examining the effect of age within the older age-group by testing its association with RSN constitution in the RSNs defined by Atlas55+. Comparison of younger and older adults showed that the SMN and VIS showed the smallest age-related differences, while the DMN, ECN and SAL showed the highest degree of age-related spatial reorganization. These findings accord with those of prior studies which have consistently reported that the functional connectivity within networks supporting higher-order functions is more affected by age than that of networks supporting lower-order functions (Mowinckel AM et al. 2012; He X *et al*. 2013; Betzel RF *et al*. 2014). Also, a recent study by Luo N, J Sui, A Abrol, J Chen, *et al*. (2020) showed more extensive age-related changes in the structural-functional coherence of the associative cortex compared with that of the unimodal cortex. The posterior medial temporal regions - which are typically identified as part of the DMN in younger adults (Buckner RL et al. 2008; Doucet G *et al*. 2011; Buckner RL and LM DiNicola 2019) - were instead assigned to the ECN in the older adults. This finding is in agreement with the notion that cognitive aging involves reorganization rather than loss of function (Reuter-Lorenz PA and C Lustig 2005; Andrews-Hanna JR et al. 2007; Yaple ZA *et al*. 2019). This regional variability of the DMN and ECN is also supported by the “the default to executive coupling” (DECHA) model of cognitive aging proposed by Turner GR and RN Spreng (2015).This model proposes a regional shift in the architecture of the DMN and ECN to support crystallized cognition in later life (Spreng RN et al. 2018). We also found large spatial differences in the SAL between the younger and older adults, with the younger adults showing a network restricted to the anterior insula bilaterally and anterior cingulate cortex only (Seeley WW *et al*. 2007). As discussed above, the origin of this large inter-individual variability is unclear. Future investigations focusing on the SAL are needed to determine whether the spatial differences may be related to analytical or true biological differences between samples.

We found a significant association of age with the functional integrity of the major RSNs in older adults, particularly in the major regions of the DMN, ECN and SAL. This finding was expected as the age range across all participants was 40 years. This was consistent with prior neuroimaging studies that described an effect of aging on both brain structure and function in populations over age 50 (Betzel RF *et al*. 2014; Chan MY et al. 2014; Luis EO et al. 2015; Damoiseaux JS 2017; Varangis E, CG Habeck, et al. 2019; Dima D et al. 2020; Frangou S et al. 2020; Luo N, J Sui, A Abrol, D Lin, *et al*. 2020). The overall negative impact of age on each network largely confirms a reduction of functional cohesiveness of the major brain networks, particularly those supporting higher-order cognitive functions (Damoiseaux JS *et al*. 2008; Mowinckel AM *et al*. 2012; He X *et al*. 2013; Betzel RF *et al*. 2014; Yaple ZA *et al*. 2019). It is interesting to note that we also found a positive association of age in cerebellar regions in both the DMN and ECN, suggesting higher cerebellar-cortical integration in these two networks with older age. Zhang H *et al*. (2017) reported similar findings of increased cerebellar-neocortical functional connectivity with age in healthy participants between the age of 12 and 79 years. This higher cerebellar-neocortical integration has also been associated with successful higher-order cognitive activity in a healthy elderly population (Luis EO *et al*. 2015). Collectively, the findings reported in the current study underscore the importance of age-adapted brain functional atlases

Moving forward, we propose Atlas55+ that has several advantages to study brain activity in late adulthood. To the best of our knowledge, Atlas55+ (a) is the only atlas based on rs-fMRI datasets from three independent cohorts including a total of 563 healthy participants above the age of 55 years; (b) includes functional partitions of the brain at the level of the large RSNs and their subdivisions; (c) accommodates differences in neuroimaging acquisition parameters (i.e., different sites, MRI scanners and acquisition sequences); and (d) is independent of sample composition. We identified 15 reproducible RSN subdivisions (Figure 4), which is in a typical range for subdivisions reported in brain functional atlases based on younger adults’ rs-fMRI data (Yeo BT *et al*. 2011; Doucet GE *et al*. 2019). We note that the subdivisions reported were those that were identified in all three older-adult cohorts, to minimize potential influences by differences in MRI acquisition parameters. Further studies are needed to confidently identify the exact causes of variation in these subdivisions between the cohorts.

While Atlas55+ offers a realistic option for standardizing the definition of RSNs for older adults, we acknowledge its limitations. First, the age cut-off of 55 years follows the criterion used in ADNI, which can be viewed as arbitrary. Second, age-related changes in RSN configuration are likely to continue throughout late life and these subtle changes may not be fully captured by Atlas55+. The option of creating brain functional atlases in older individuals to cover specific and smaller time periods (i.e., for every decade past the 55^th^ year) is challenging at present as high-quality rs-fMRI datasets are scarce in older participants, particularly in advanced old age. The Human Connectome Project-Aging dataset may help achieve this goal when it will be fully released (https://www.humanconnectome.org/study/hcp-lifespan-aging). Third, we constructed Atlas55+ using a volumetric-based (ICA) rather than a surface-based approach. We previously demonstrated that MICCA provided reliable networks (Naveau M *et al*. 2012), and the analytic approach does not significantly alter the reproducibility of the RSNs as long as they are derived from a sample of at least 100 individuals (Doucet GE *et al*. 2019), which is the case for each cohort analyzed in the current study. Therefore, it is unlikely that this choice strongly influenced the spatial definition of the RSNs. Fourth, we focused on the five major RSNs (DMN, ECN, SAL, SMN, VIS) and their subdivisions as these are the most reproducible networks in the neuroimaging literature (van den Heuvel MP and HE Hulshoff Pol 2010; Doucet GE *et al*. 2019; Elliott ML *et al*. 2019). As aging is associated with reduced brain modularity (Meunier D *et al*. 2009; Damoiseaux JS 2017; Varangis E, Q Razlighi, *et al*. 2019), we cannot exclude the possibility that the clustering approach did not influence the current findings (Abou Elseoud A *et al*. 2011). However, each of these five RSNs have been consistently identified in aging studies (Damoiseaux JS *et al*. 2008; Mowinckel AM *et al*. 2012; Betzel RF *et al*. 2014; Sala-Llonch R et al. 2015; Varangis E, CG Habeck, *et al*. 2019), suggesting that their existence in older individuals is indisputable. Lastly, this study specifically focused on the spatial characteristics of the RSNs; other characteristics such as their dynamics or the association with cognitive variability in late adulthood were not explored because the necessary data were not available. In particular, it will be important to investigate the cognitive impact on the spatial definition of the Atlas55+ RSNs, in cognitively impaired populations. In the current study, all individuals were identified as healthy and showed very low variability in their general functioning (Mini-mental state exam score: mean (std): 28.6 (1.5), across the CamCAN55+ and ADNI cohorts) which prevented further analyses to identify spatial variability in the RSNs as a function of cognitive ability.

## Conclusion

We presented a quantitative comparison of the spatial composition of the major RSNs and their reliable functional subdivisions derived from healthy adults between the ages of 55 and 95 years old and younger adults aged 18-35 years. We also identified the spatial distribution of age-related changes in RSN definition. Our results confirm the importance of age-appropriate brain functional atlases for studies investigating brain mechanisms related to aging. We propose that Atlas55+ can provide a reproducible template of the major RSNs for late adulthood as it is publicly available (Website-to-be-provided upon publication).

## Acknowledgments

This work was supported by the National Institute of Aging (R03AG064001 to G.E.D); the National Institute of General Medical Sciences (P20GM130447 to G.E.D.); the National Institute of Mental Health (R01MH113619 to S.F); the National Institute of Biomedical Imaging and Bioengineering (U54 EB020403 to P.M.T.); the French government agency ANR (LABCOM Ginesislab, ANR 16-LCV2-0006-01 to M.J.). LL is supported by a fellowship from the French Alternative Energies and Atomic Energy Commission (CEA). Data collection and sharing for this project was funded by the Alzheimer’s Disease Neuroimaging Initiative (ADNI) (National Institutes of Health Grant U01 AG024904) and DOD ADNI (Department of Defense award number W81XWH-12-2-0012). ADNI is funded by the National Institute on Aging, the National Institute of Biomedical Imaging and Bioengineering, and through generous contributions from the following: AbbVie, Alzheimer’s Association; Alzheimer’s Drug Discovery Foundation; Araclon Biotech; BioClinica, Inc.; Biogen; Bristol-Myers Squibb Company; CereSpir, Inc.; Cogstate; Eisai Inc.; Elan Pharmaceuticals, Inc.; Eli Lilly and Company; EuroImmun; F. Hoffmann-La Roche Ltd and its affiliated company Genentech, Inc.; Fujirebio; GE Healthcare; IXICO Ltd.; Janssen Alzheimer Immunotherapy Research & Development, LLC.; Johnson & Johnson Pharmaceutical Research & Development LLC.; Lumosity; Lundbeck; Merck & Co., Inc.; Meso Scale Diagnostics, LLC.; NeuroRx Research; Neurotrack Technologies; Novartis Pharmaceuticals Corporation; Pfizer Inc.; Piramal Imaging; Servier; Takeda Pharmaceutical Company; and Transition Therapeutics. The Canadian Institutes of Health Research is providing funds to support ADNI clinical sites in Canada. Private sector contributions are facilitated by the Foundation for the National Institutes of Health (www.fnih.org). The grantee organization is the Northern California Institute for Research and Education, and the study is coordinated by the Alzheimer’s Therapeutic Research Institute at the University of Southern California. ADNI data are disseminated by the Laboratory for Neuro Imaging at the University of Southern California.

## Data and code availability

The datasets used in the current study are freely available or upon request (links provided in supplementary material). The RSNs extracted for each cohort and from Atlas55+ are publicly available (Website-to-be-provided upon publication) and upon request to the corresponding author.

