## Supplementary Material for "Atlas55+: Brain Functional Atlas of Resting-state Networks for Late Adulthood"

**Method**

1. **Cohort Description**

The CamCAN cohort is available upon request ([www.mrccbu.cam.ac.uk/datasets/camcan](http://www.mrccbu.cam.ac.uk/datasets/camcan)) and includes a total of 652 individuals (330 Females, 18-88 years old), with 326 between the age of 55 and 88 years (Shafto MA et al. 2014; Taylor JR et al. 2017). Among this subsample, we excluded 74 participants because they reported psychiatric disorders, history of head injury with loss of consciousness, cancer, or epilepsy, and 2 participants that did not have both rs-fMRI and structural MRI data available, from further analyses.

The Southwest University Adult Lifespan Dataset (SALD) cohort is publicly available through the International Data-sharing Initiative (INDI, <http://fcon_1000.projects.nitrc.org/indi/retro/sald.html>) and includes a total of 494 healthy individuals (308 Females, 19-80 years old), with 190 between the age of 55 and 80 years (Wei D et al. 2018). The exclusion criteria for this dataset included: (1) MRI related exclusion criteria, which included claustrophobia, metallic implants, Meniere’s Syndrome and a history of fainting within the previous 6 months; (2) current psychiatric disorders and neurological disorders; (3) use of psychiatric drugs within the three months prior to scanning; (4) pregnancy; or (5) a history of head trauma (Wei D *et al.* 2018).

The Alzheimer’s Disease Neuroimaging Initiative (ADNI) cohort is available upon request (<http://adni.loni.usc.edu/>) and is an on-going project including the collection of longitudinal behavioral and neuroimaging data from 819 individuals enrolled between the age of 55 and 90 years (healthy participants, patients with mild cognitive impairment (MCI) or Alzheimer’s Disease (AD) (Jack CR, Jr. et al. 2008; Petersen RC et al. 2010). The ADNI was launched in 2003 as a public-private partnership, led by Principal Investigator Michael W. Weiner, MD. The primary goal of ADNI has been to test whether serial MRI, PET, other biological markers, and clinical and neuropsychological assessment can be combined to measure the progression of mild cognitive MCI and early AD. As of November 2019, both resting-state fMRI and structural MRI data from 136 healthy participants were available from Waves 2 and 3 of the project and were considered for the current study.

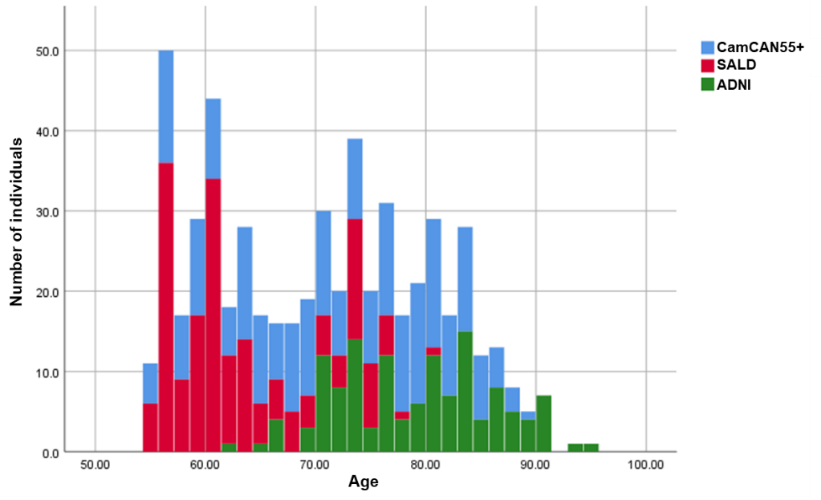

**Supplementary Figure S1: Stacked distribution of the age across the three cohorts.**

2. **Acquisition and pre-processing of the resting-state fMRI data**

**2.1. Acquisition**

In the CamCAN cohort, rs-fMRI data were acquired while participants rest with their eyes closed on a 3T Siemens TIM Trio scanner with a 32-channel head coil with the following acquisition parameters: TR/TE= 1970/30 msec, 32 axial slices, flip angle =78 degrees; FOV =192 mm×192 mm; voxel-size =3 mm×3 mm×4.44 mm, acquisition time=8min 40sec, number of volumes: 261. More information on the MRI sequences can be found in Taylor JR *et al.* (2017).

In the SALD cohort, rs-fMRI data were acquired while participants rest with their eyes closed on a 3T Siemens Trio scanner, using the following acquisition parameters (Wei D *et al.* 2018): TR/TE= 2000/30 msec, 32 axial slices, flip angle=90 degrees; FOV=220 mm×220 mm; voxel-size =3.4 mm×3.4 mm×4 mm, acquisition time=8min, number of volumes: 242. More information on the MRI sequences can be found in Wei D *et al.* (2018).

In the ADNI cohort, rs-fMRI data were acquired while participants rest with their eyes open, according to the ADNI acquisition protocol (Jack CR, Jr. *et al.* 2008), over 43 sites, using 3T scanners (General Electric (GE), Philips or Siemens). All rs-fMRI data were collected using a TR=3000 sec, with 140 to 200 volumes (detail in <http://adni.loni.usc.edu/methods/documents/>). When multiple resting-state fMRI datasets were available for a participant, the first one was selected for analyses.

**2.2. Preprocessing**

Regardless of the cohort, resting-state fMRI data were preprocessed using SPM12 and the DPABI Toolbox (Yan CG et al. 2016). The preprocessing procedures for the resting-state fMRI datasets included removal of the first 3 volumes, motion correction to the first volume with rigid-body alignment; coregistration between the functional scans and the anatomical T1-weighted scan; spatial normalization of the functional images into Montreal Neurological Institute (MNI) stereotaxic standard space; spatial smoothing within the functional mask with a 6-mm at full-width at half-maximum Gaussian kernel; wavelet despiking (Patel AX et al. 2014); linear detrending; and regression of motion parameters and their derivatives (24-parameter model) (Friston KJ et al. 1996), as well as white matter (WM), CSF time series. The WM and CSF signals were computed using a component based noise reduction method (CompCor, 5 principal components) (Behzadi Y et al. 2007). Lastly, bandpass filtering was applied at [0.01-0.1] Hz (Cordes D et al. 2001).

**2.3.** **Data Quality Assurance**

Across the three late-life cohorts, we excluded a total of 11 individuals for having excessive head movement based on maximum transient (volume-to-volume) head motion above 2 mm translation or 1 degree rotation, leading to a final sample of: 246 for CamCAN, 185 for SALD, and 132 for ADNI. Further, we computed the framewise displacement (FD) in the rs-fMRI datasets using a Matlab function freely available

(https://github.com/spunt/bspm/blob/master/thirdparty/bramila/bramila_framewiseDisplacement.m) which follows the formula provided by Power JD et al. (2012). After removal of these datasets, the mean FD did not correlate with age (r=0.08).

In the CamCAN35- sample (individuals of 35 years and youngers), we excluded two individuals for having excessive head movement based on maximum transient (volume-to-volume) head motion above 2 mm translation or 1 degree rotation.

3. **Multi-scale clustering of individual component algorithm (MICCA)**

**3.1. Group-level components**

This step aimed to identify statistically reliable brain networks using a process validated by Naveau et al (2012).

First, for each individual within each cohort, single subject independent component analyses (ICAs) were conducted 20 times with random initialization using the Multivariate Exploratory Linear Optimized Decomposition into Independent Components (MELODIC) software, version 3.15, included in the FMRIB Software Library (FSL) v6.0.3 (http://www.fmrib.ox.ac.uk/fsl) (Smith SM et al. 2004). The number of independent components (ICs) was estimated by Laplace approximation (Minka T 2000). A symmetric approach of the FastICA algorithm (Hyvarinen A 1999) was used to compute the ICAs. Second, for each repetition, we used the multi-scale clustering of individual component algorithm (MICCA) (Naveau M et al. 2012) to classify ICs in *N* groups. Note that the number of groups was automatically estimated by the algorithm. Third, the Icasso algorithm (Himberg J et al. 2004) was used to select the groups of reproducible ICs (i.e., groups with ICs that were present in at least 50% of the 20 repetitions), which we identified as “group-level components” (Salman MS et al. 2019). Third, for each group-level component, a voxel-wise t-score map was computed and thresholded using a mixture model (p>0.95) (Beckmann CF and SM Smith 2004). Lastly, we discarded the group-level components if their spatial map: (i) mainly covered non-gray matter (i.e., CSF and WM), or (ii) included regions with strong signal attenuation due to susceptibility artefacts (i.e., lower frontal and lower temporal regions) (Supplementary Figures S2B-S4B).

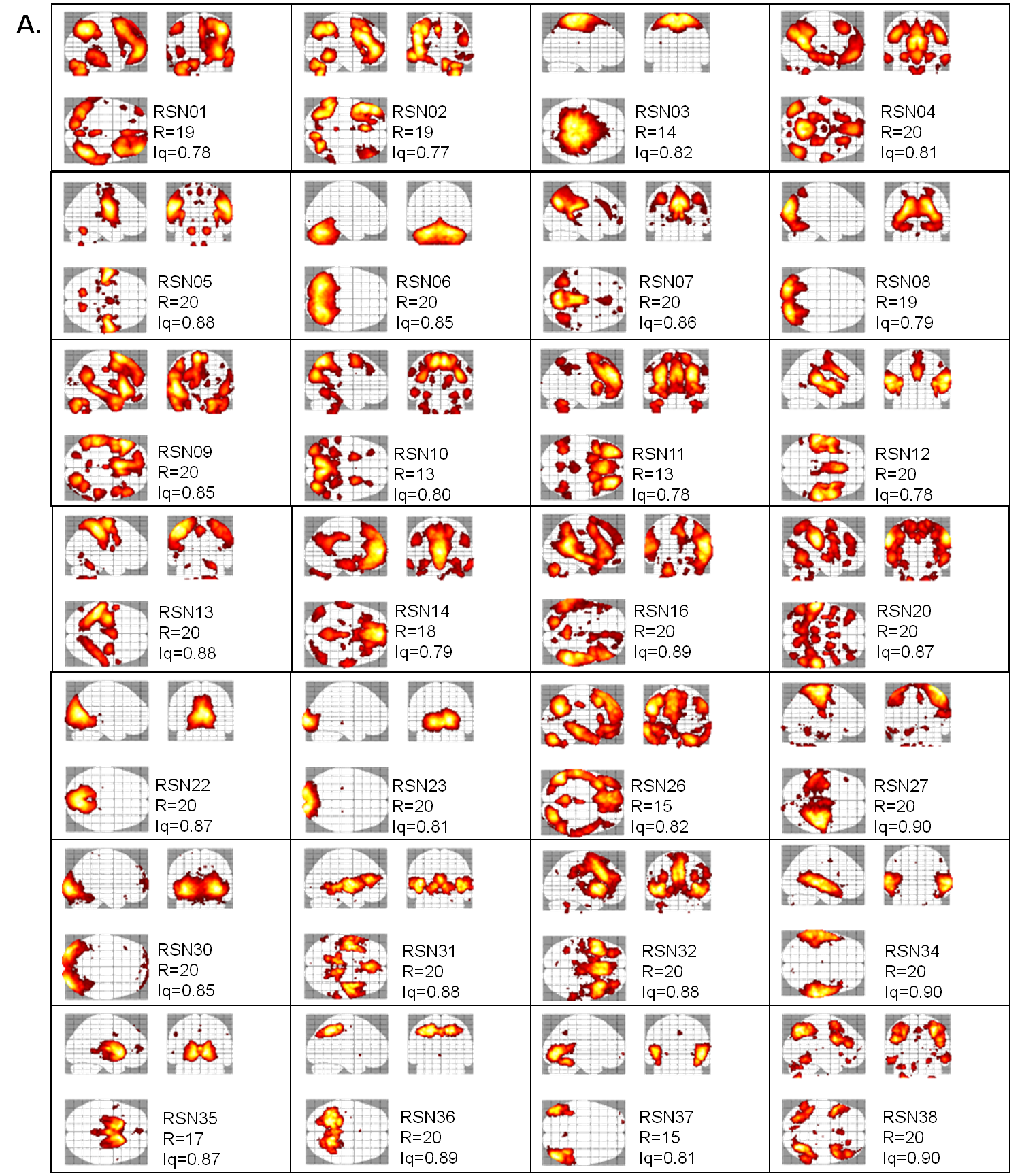

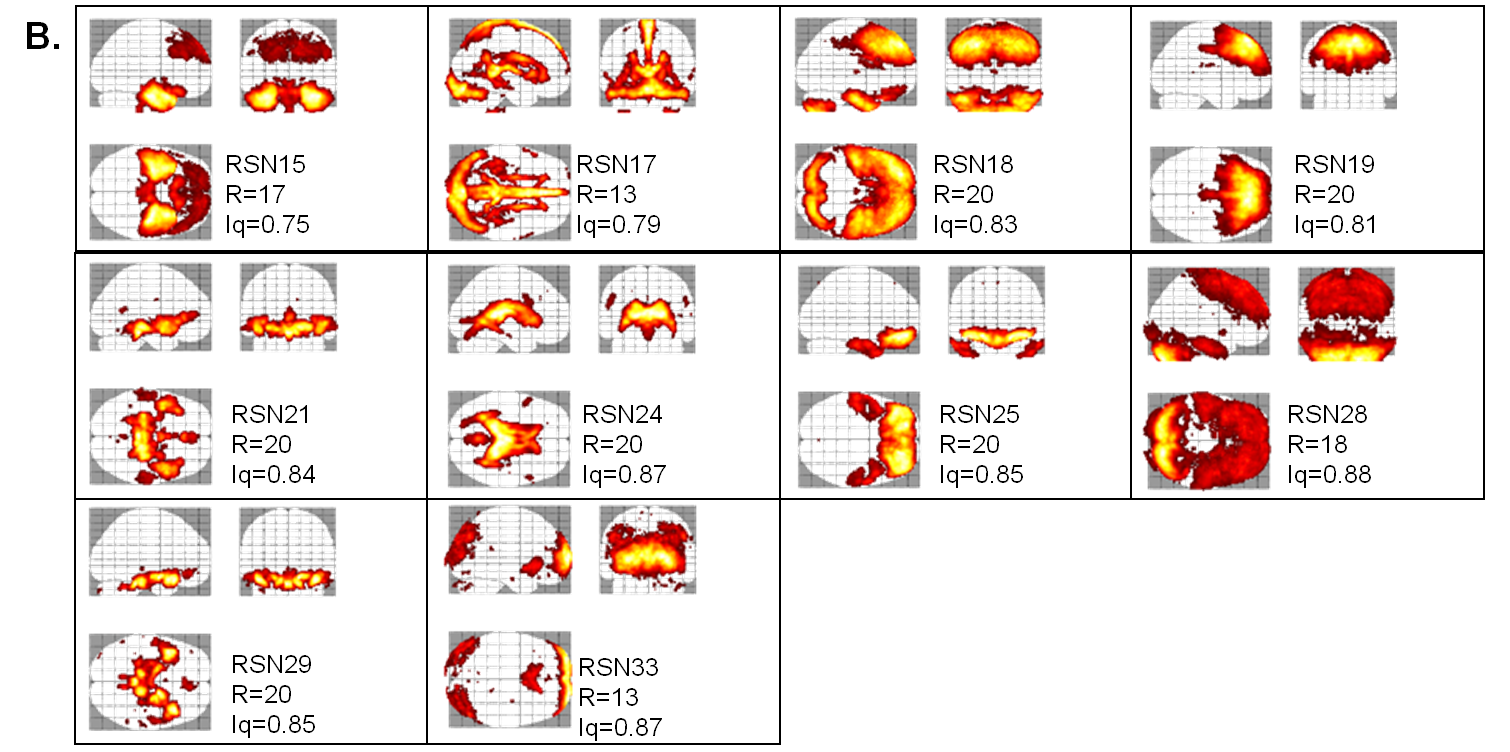

**Supplementary Figure S2: Spatial maps of the group-level components in the CamCAN55+ cohort.** (A) Included group-level components; (B) excluded group-level components. R= Number of ICA repetitions that identified the group-level component. Iq= Quality Index, based on the Icasso algorithm (Himberg J *et al.* 2004).

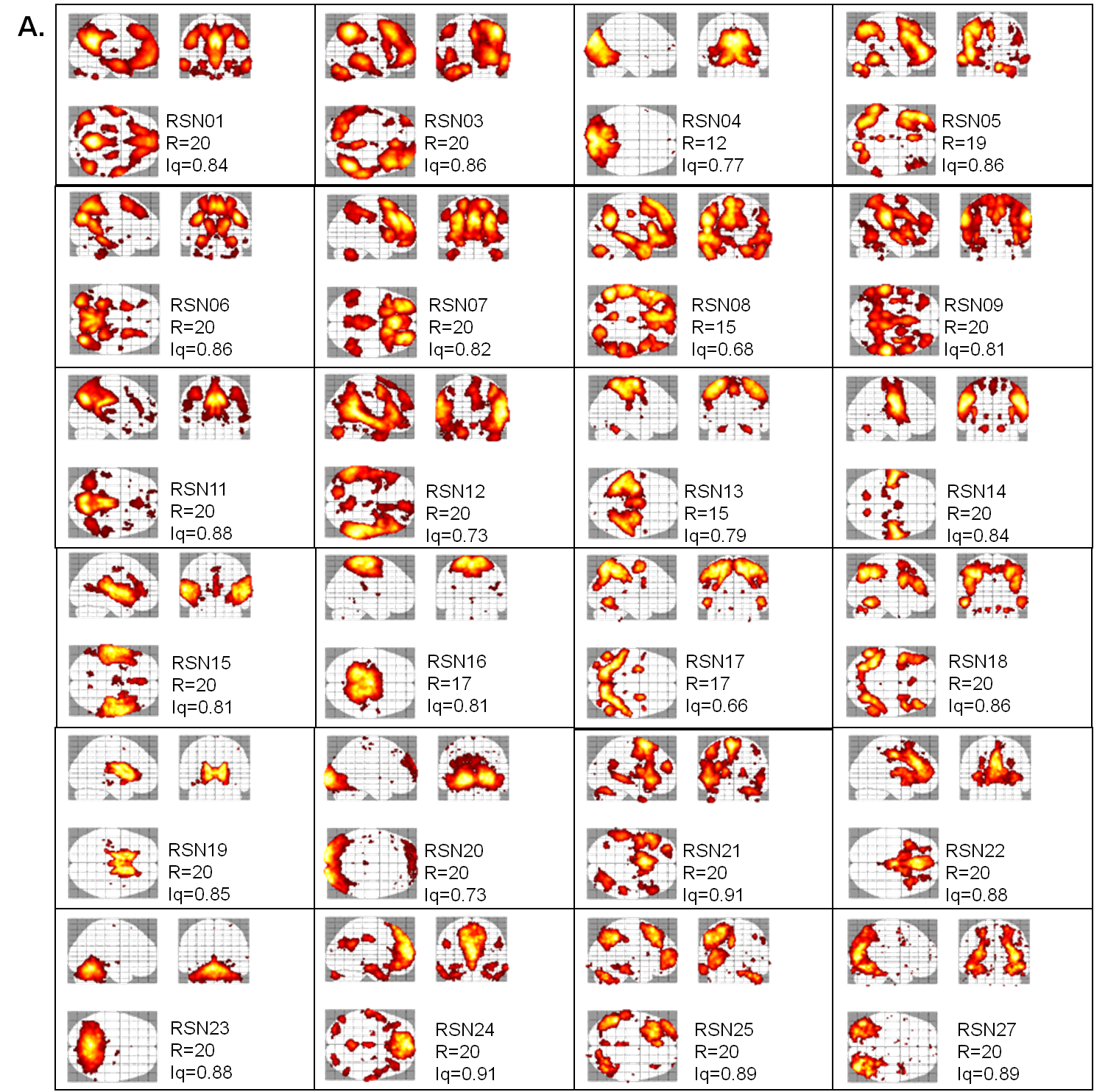

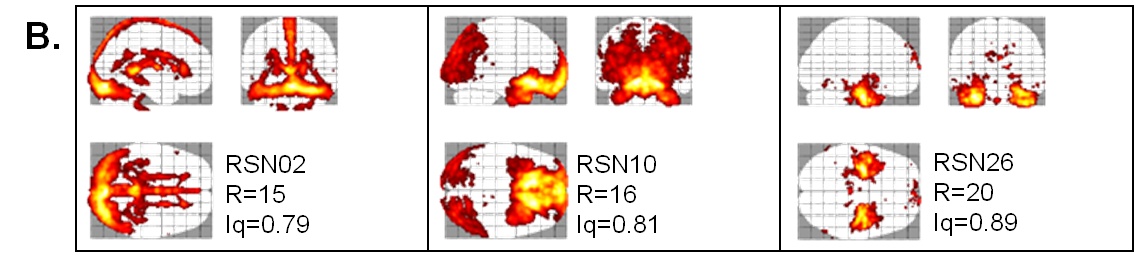

**Supplementary Figure S3: Spatial maps of the group-level components in the SALD cohort.** (A) Included group-level components; (B) excluded group-level components. R= Number of ICA repetitions that identified the RSN. Iq= Quality Index, based on the Icasso algorithm (Himberg J *et al.* 2004).

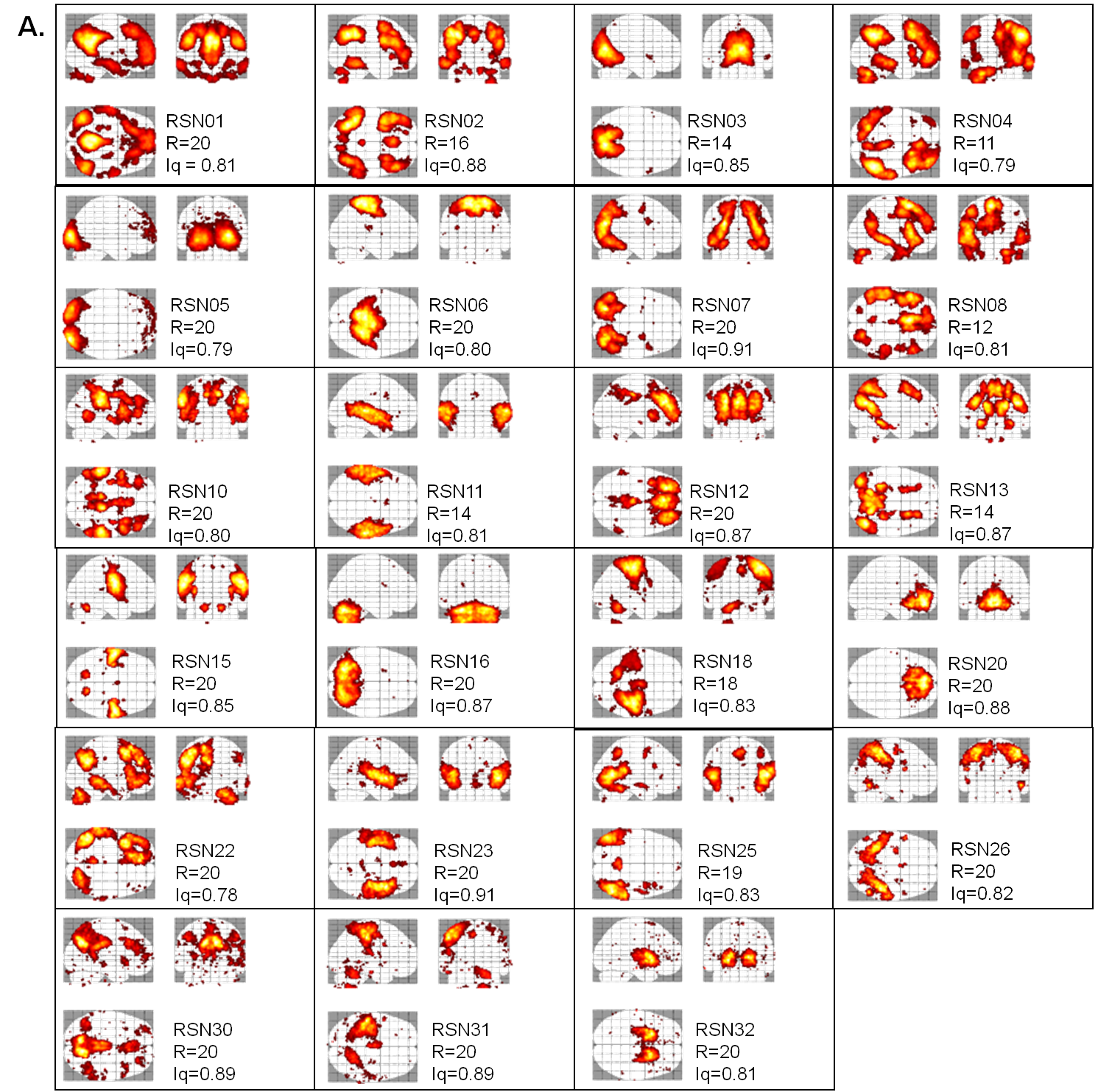

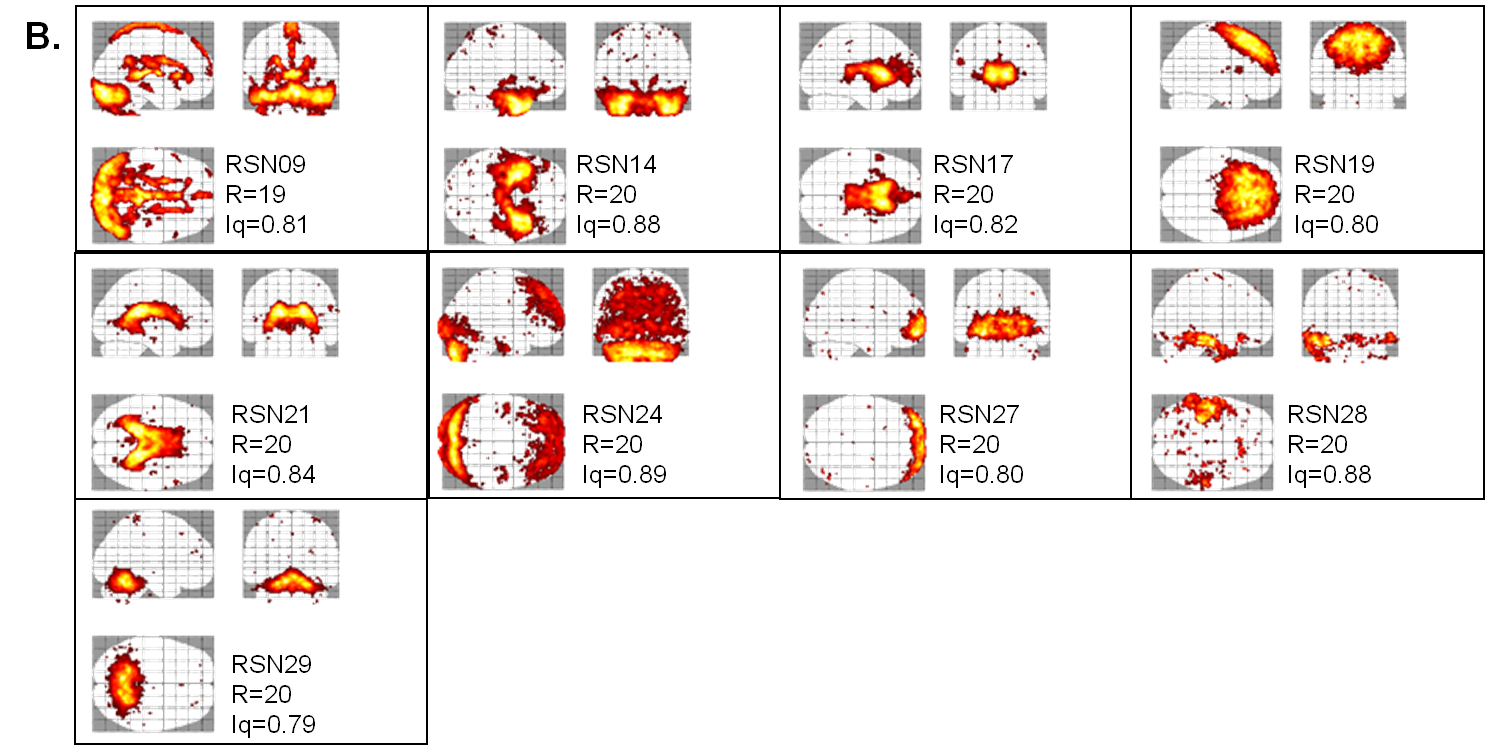

**Supplementary Figure S4: Spatial maps of the group-level** **components in the ADNI cohort.** (A) Included group-level components; (B) excluded group-level components. R= Number of ICA repetitions that identified the group-level components. Iq= Quality Index, based on the Icasso algorithm (Himberg J *et al.* 2004).

**3.4. Functional Reliability of the group-level components: Computation of the Q index**

The reliability Q score represents the relative position of the network *k* across the individual dendrograms, relative to the group (average) dendrogram (Supplementary Figure S5). In summary, the first step of the process involves the build of a group (average across individuals of a single cohort) and single-subject dendrograms, following a hierarchical ascendant clustering conducted on the functional connectivity matrices of dimension K*K (where K is the number of components) (Doucet et al. 2011) (Supplementary Figure S5A). The structure of a dendrogram can be then defined as a partition *P^j^* of K vectors, with j=1 to K. An individual partition for subject s is defined as: *P^j^_s_* and average partition is defined as *P^j^_m_*. Each vector is defined as follows: the *k^th^* component is equal to 1 if the network *k* is in the same cluster than the network *j*; and 0 if it is not (Supplementary Figure S5B). The second step involves the comparison of each individual partition *P^j^_s_* to the average partition *P^j^_m_*, by computing the Sorensen-Dice (D) indices. Lastly, we aimed to quantify the proportion of individual partitions that were the most similar to the average partition, for each network. We therefore identified the individual partitions with the highest D that were different from 1 and referred them as the "best alternative partitions". In this context, the Q score for network *k* is defined as follows (Supplementary Figure S5C):

$Q_{k}=\frac{\sum_{j=1}^{K} N_{k}^{j}}{\sum_{j=1}^{K} N^{j}}$,

where *N^j^_k_* as the number of subjects expressing the best alternative partition containing the network *k* for a given average partition *P^j^_m_,* and *N^j^* as the number of subjects expressing the best overall alternative partition for a given average partition *P^j^_m_*.

We conducted Tukey's fences tests to identify and discard these highly unreliable group-level components. Group-level components with a Q score greater than 1.5 times the interquartile range of the average Q of all components in a cohort were considered highly unreliable. We did not identify any outliers (Supplementary Figure S6).

More detail on the description and validation of the Q index can be found in Labache L et al. (2020).

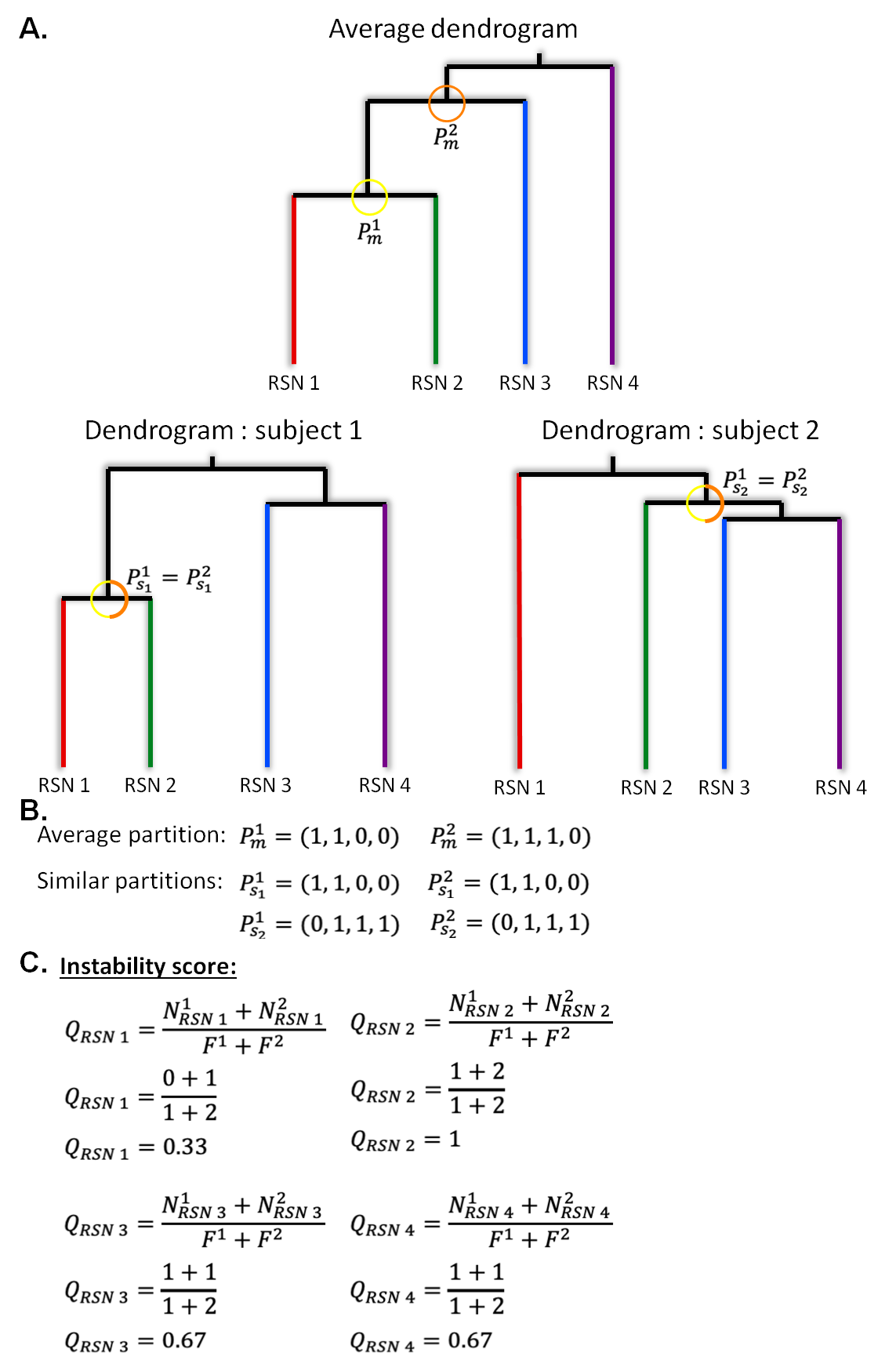

**Supplementary Figure S5: Simplified schematic of the process to compute the instability Q index for each group-level** **component.** This example is based on 2 subjects (s1 and s2), 4 group-level components (or resting-state networks (RSNs)). This index is based on resting-state functional connectivity between the pairs of networks.

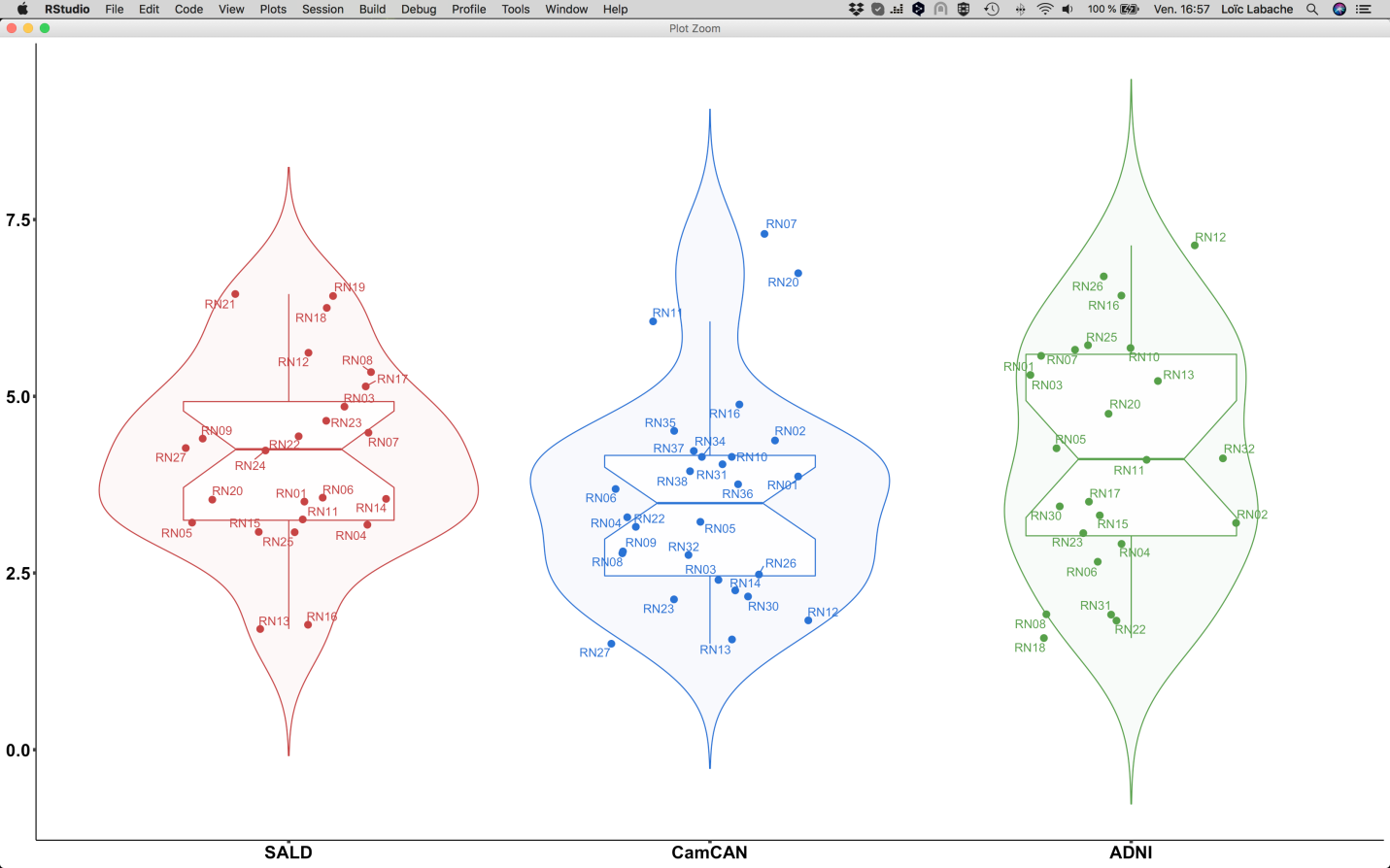

**Supplementary Figure S6: Boxplots of the instability Q scores for each group-level component identified in the SALD, CamCAN55+ and ADNI cohorts.**

**3.5. Maps of spatial overlap between group-level** **components**

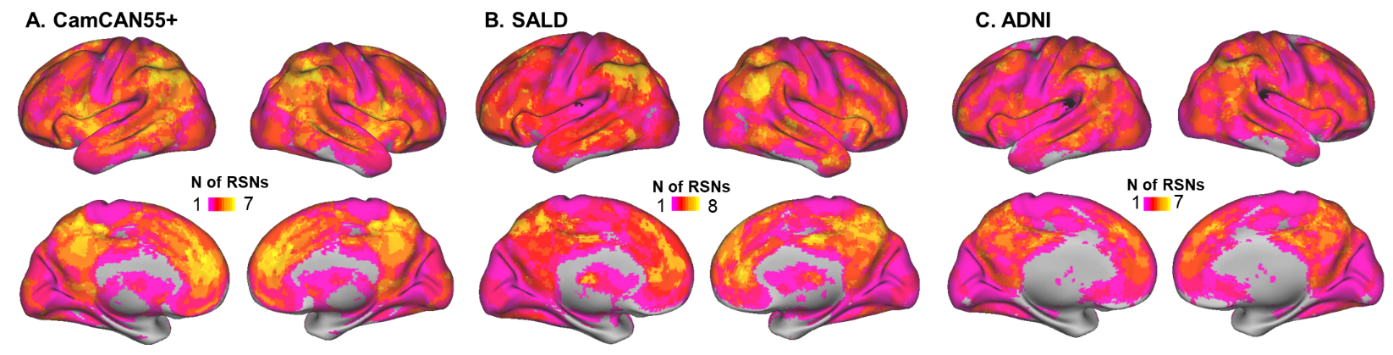

**Supplementary Figure S7: Voxelwise map for the spatial overlap across the group-level components, in the CamCAN55+ (A), SALD (B) and ADNI (C) Cohorts.**

**3.6. Non-overlapping group-level components**

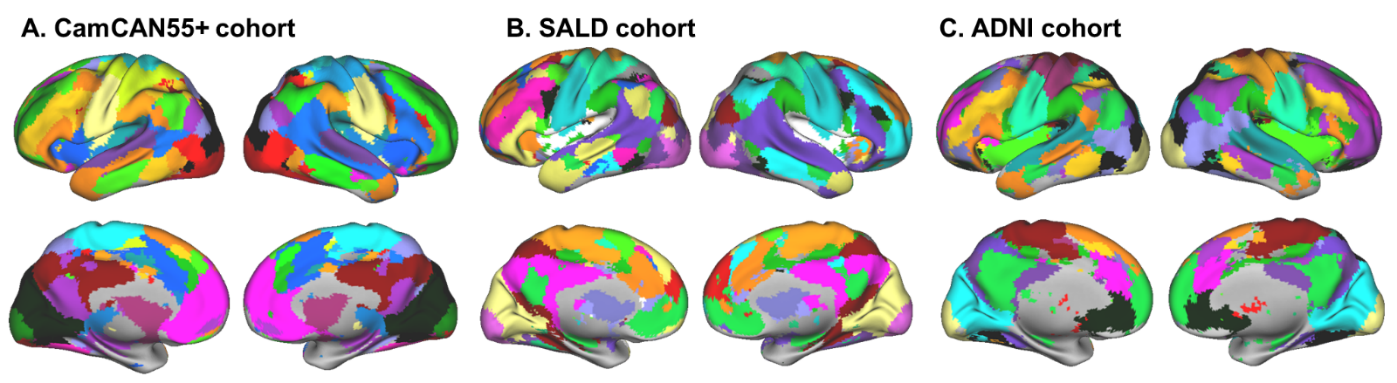

**Supplementary Figure S8: Spatial maps of the non-overlapping group-level components, in the CamCAN55+ (A), SALD (B) and ADNI (C) Samples.** One color per network.

4. **Construction of the 5 major RSNs in each late-life cohort**

We assigned each RSN to one of the major RSNs (DMN, ECN, SAL, VIS and SMN) identified in the Consensual Atlas of REsting-state Networks (CAREN) (Doucet GE et al. 2019). We used the Dice’s coefficient to quantify the degree of spatial overlap between the group-level components and the CAREN RSNs and assigned each group-level component to the RSN with the largest overlap following a winner-takes-all rule (Supplementary Table S1). We were thus able to construct a spatial map for each RSN. We repeated this operation in each cohort independently.

| **Supplementary Table S1: Spatial Matching Between non-overlapping group-level components and CAREN-RSNs.** | | | | |
| --- | --- | --- | --- | --- |
| **SensoriMotor Network (SMN)** | **Visual Network (VIS)** | **Salience Network (SAL)** | **Default Mode Network (DMN)** | **Executive Central Network (ECN)** |
| **CamCAN55+** | | | | |
| 3, 5, 13, 27, 34 | 8, 22, 23, 30, 37 | 12, 20, 32 | 4, 7, 9, 14, 16, 26 | 1, 2, 10, 11, 36, 38 |
| **SALD** | | | | |
| 13,14,15,16 | 4,20,27 | 7, 9, 21, 22 | 1, 8, 11, 12, 24, 25 | 3, 5, 6, 17, 18 |
| **ADNI** | | | | |
| 6, 11, 15, 18, 31 | 3, 5, 25 | 10, 12, 23 | 1, 8, 20, 22 | 2, 4, 13, 26, 30 |
| Number refers to the group-level components number identified by MICCA, and described in Supplementary Figures S2-S4. Group-level components mostly covering cerebellum and subcortical regions were not included. | | | | |

**4.1. Reproducibility of the spatial definition of the five RSNs**

In order to ensure that the results were not dependent on the template used, we also used the 7-network atlas created by Yeo BT et al. (2011) instead of the CAREN to do the assignment. In order to match the 5 RSNs from CAREN, we excluded the dorsal attentional and the affective networks from the Yeo Atlas from the matching procedure. We found very high similarity with regards to the spatial definition of the 5 RSNs as the average spatial overlap between same-label networks were 86% across the three cohorts (CamCAN: 86% (sd=15%), SALD: 85% (19%), ADNI: 86% (9%)) (Supplementary Figure S10).

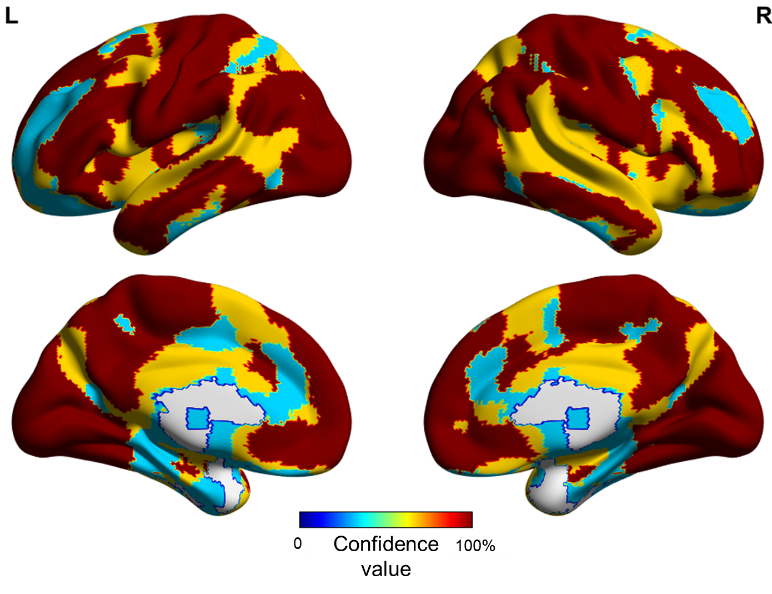

**Supplementary Figure S9: Confidence map of the reproducibility of the RSNs across the three late-life cohorts.** This measure quantifies the probability that a voxel is assigned to the same-label RSN in each of the 3 cohorts. Scores of 100% indicate complete consistency of network assignment across the 3 cohorts; Scores closer to 0 indicate lower confidence in network assignment across the cohorts.

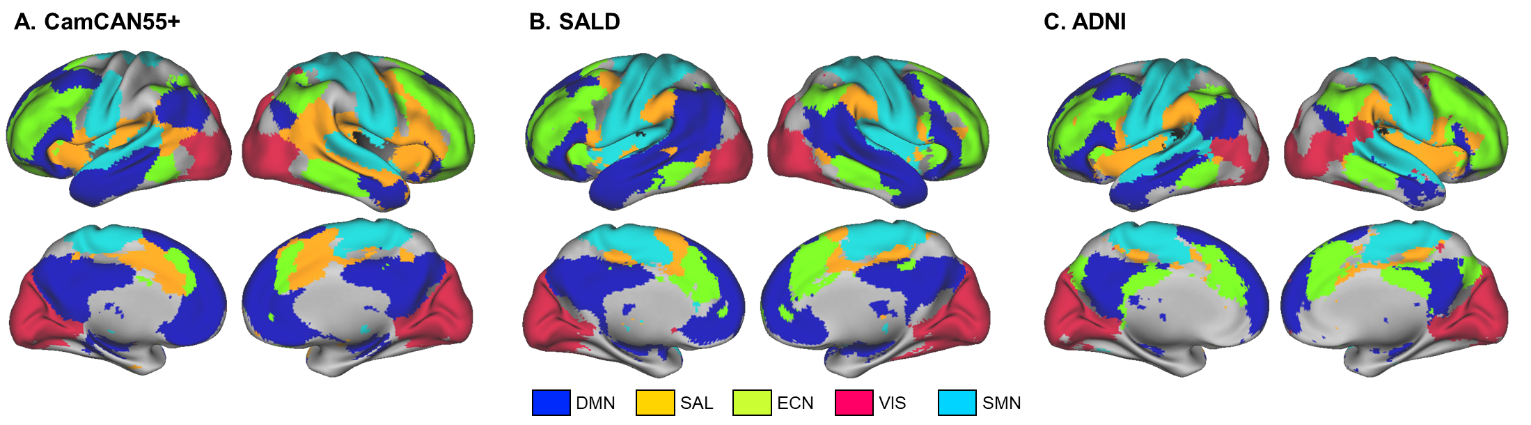

**Supplementary Figure S10: Major RSNs based on the Yeo's Atlas Assignment.**

4.2. **Subdivisions of the major RSNs in Atlas55+**

With Atlas55+, the brain was examined for a coarse solution that organized the cortex into 5 RSNs. We further report a ﬁner solution that identiﬁed 15 subnetworks that were present across the three late-life cohorts.

Within each major RSN, we computed the Dice coefficients between the group-level components assigned to the RSN across the three cohorts (Supplementary Table S1). Using the degree of spatial overlap across each pair of components, we further constructed a dendrogram in order to identify clusters of three independent group-level components with high spatial overlap, with one component from each cohort (Supplementary Figure S11). Lastly, for each of these clusters, we created a spatial map representing a subnetwork of the RSN. A voxel was identified as part of the subnetwork if it was found in at least two out of the three components. We chose to focus on clusters with three independent components because this suggests that these components can be found in independent cohorts and therefore are reliable and less influenced by acquisition parameters.

We revealed a total of 15 RSNs (Figure 5, Supplementary Table S3), which is consistent with other atlases such as the one by Yeo et al. (2011) which described a 17-network brain decomposition in younger adults. We chose to focus on subdivisions that were reliably identified in all three late-life cohorts. Less reproducible RSN subdivisions were also present as they were only identified across two cohorts. Because of the discrepancies in demographics, or MRI acquisition parameters between the cohorts, it is not possible to provide a clear explanation on the origins of these subdivisions differences.

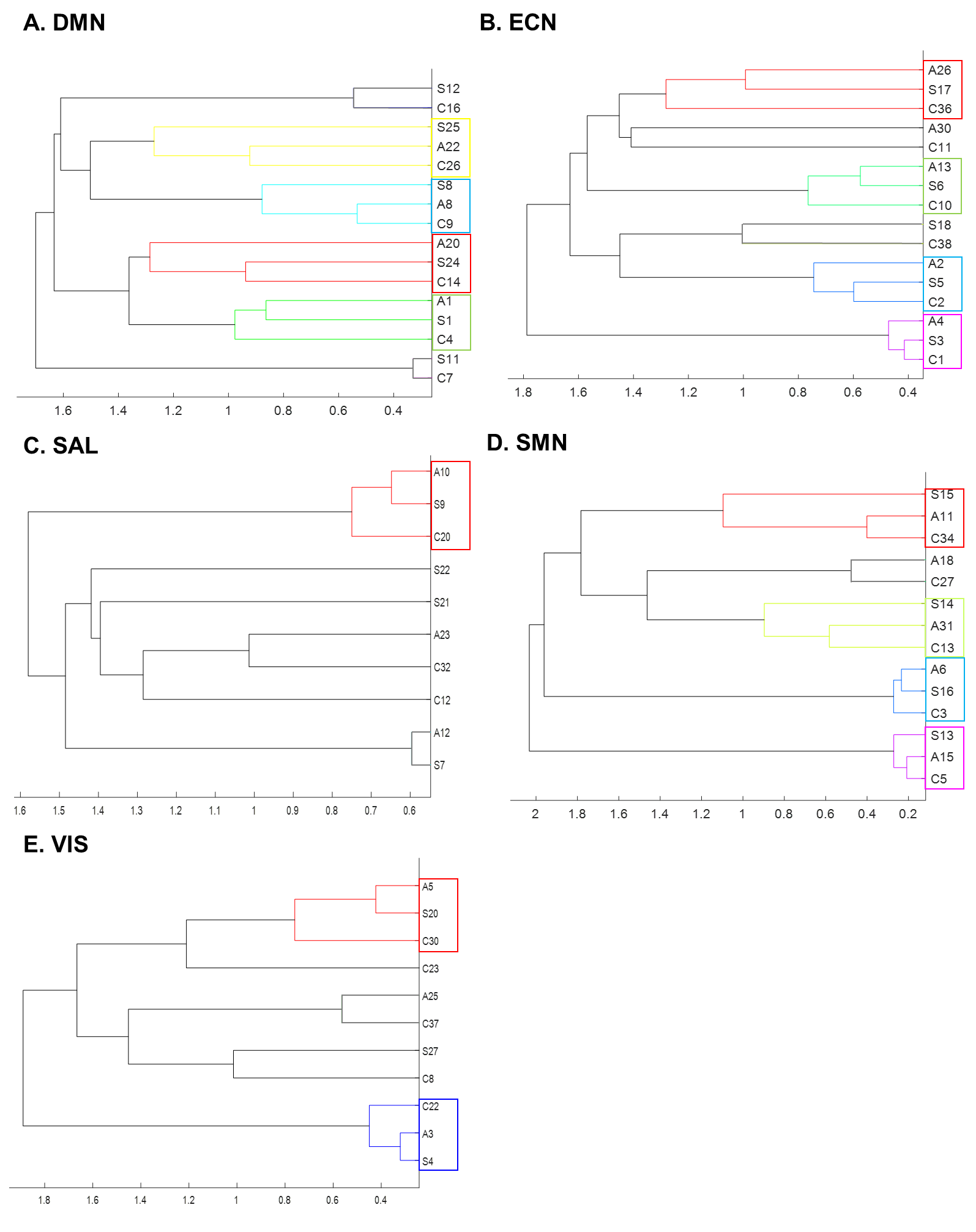

**Supplementary Figure S11: Dendrograms based on the spatial overlap (Dice coefficients) between group-based components, for each major RSN.** Each component’s assignment is listed in Supplementary Table S1; where the letter refers to the cohort (“A” refers to group-based components from ADNI, “S” refers to group-based components from SALD, “C” refers to group-based components from CamCAN55+), and the number refers to the number of the component assigned to a specific RSN. Each cluster of three independent components is highlighted in color and further identified as a RSN subdivision. The associated spatial map of these RSN subdivisions is shown in Figure 4 of the main text, with the same color code and anatomically described in Supplementary Table S3.

5. **Spatial Definition of the RSNs in mid-adulthood (from CamCAN35- cohort)**

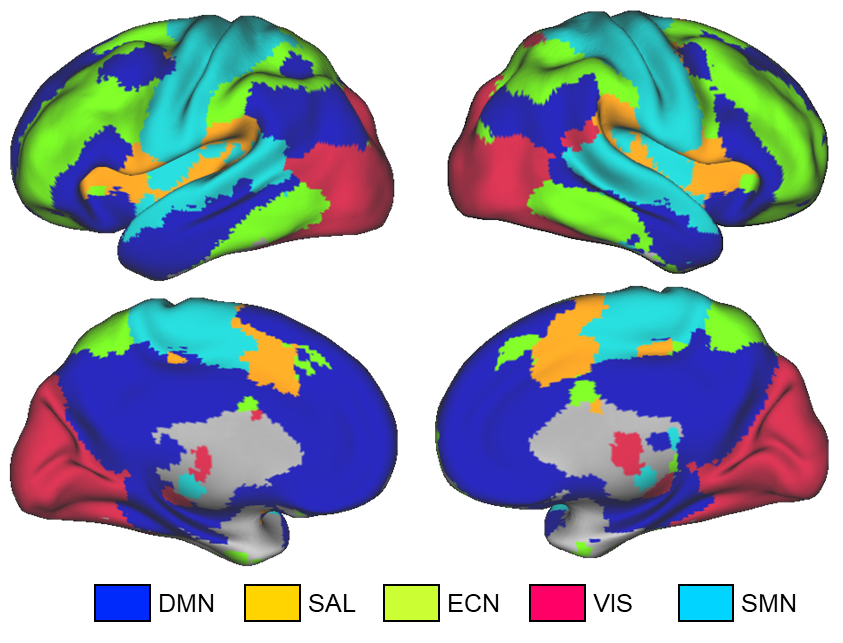

**Supplementary Figure S12: Spatial map of the major resting-state networks in the CamCAN35- cohort (individuals aged 18-35 years).**

6. **Atlas55+**

**6.1. Anatomical definition of the Atlas55+ RSNs**

| **Supplementary Table S2: Anatomical Description of the 5 RSNs of Atlas55+.** | | | | | | | |
| --- | --- | --- | --- | --- | --- | --- | --- |
| **Regions** | **Hemisphere** | **p(FWE-corr)** | **N of voxels** | **T** | **x** | **y** | **z** |
| **Default Mode Network** | | | | | | | |
| Posterior Cingulate Cortex | L | <1E-15 | 7243 | 56.04 | -6 | -52 | 34 |
| Posterior Cingulate Cortex | R |  |  | 52.51 | 4 | -54 | 32 |
| Middle Temporal Gyrus | L | <1E-15 | 21877 | 52.82 | -60 | -16 | -10 |
| Middle Temporal Gyrus | L |  |  | 50.49 | -62 | -30 | -4 |
| Angular Gyrus | L |  |  | 50.08 | -50 | -64 | 30 |
| Middle Temporal Gyrus | R | <1E-15 | 9410 | 49.37 | 60 | -2 | -16 |
| Middle Temporal Pole | R |  |  | 46.52 | 54 | 8 | -22 |
| Cerebellum Crus I | L | <1E-15 | 478 | 35.16 | -26 | -78 | -32 |
| Cerebellum Crus I | R | <1E-15 | 1359 | 32.17 | 24 | -80 | -26 |
| Cerebellum Crus I | R |  |  | 26.08 | 30 | -78 | -32 |
| Cerebellum Area 9 | R | <1E-15 | 145 | 16.27 | 6 | -52 | -40 |
| Cerebellum Area 9 | L |  |  | 14.69 | -6 | -52 | -40 |
| Middle Frontal Gyrus | R | <1E-15 | 147 | 13.87 | 40 | 10 | 48 |
| Middle Frontal Gyrus | R |  |  | 10.42 | 34 | 18 | 36 |
| **Executive Control Network** | | | | | | | |
| Angular Gyrus | R | <1E-15 | 13232 | 55.47 | 36 | -64 | 48 |
| Inferior Parietal Gyrus | R |  |  | 50.21 | 46 | -46 | 52 |
| Inferior Temporal Gyrus | R | <1E-15 | 2101 | 53.53 | 58 | -48 | -10 |
| Inferior Temporal Gyrus | L | <1E-15 | 1376 | 52.22 | -56 | -56 | -10 |
| Inferior Temporal Gyrus | L |  |  | 10.87 | -60 | -18 | -28 |
| Cerebellum Crus I | L | <1E-15 | 2076 | 46.30 | -10 | -74 | -26 |
| Cerebellum Area 6 | L |  |  | 40.88 | -28 | -62 | -30 |
| Middle Frontal Gyrus | R | <1E-15 | 7768 | 42.35 | 30 | 12 | 60 |
| Inferior Frontal Gyrus | R |  |  | 38.21 | 46 | 38 | 26 |
| Inferior Frontal Gyrus | R |  |  | 33.36 | 48 | 18 | 40 |
| Middle Frontal Gyrus | L | <1E-15 | 4657 | 39.42 | -28 | 8 | 62 |
| Middle Frontal Gyrus | L |  |  | 34.76 | -46 | 32 | 30 |
| Cerebellum Crus I | R | <1E-15 | 325 | 33.66 | 10 | -76 | -26 |
| Cerebellum Area 6 | R | <1E-15 | 253 | 32.33 | 28 | -62 | -30 |
| Cerebellum Crus I | R |  |  | 27.85 | 36 | -68 | -28 |
| Cerebellum Area 6 | L | 2.22E-15 | 95 | 18.24 | -34 | -40 | -36 |
| Cerebellum Area 10 | R | <1E-15 | 159 | 16.79 | 30 | -32 | -38 |
| Cerebellum Area 10 | R |  |  | 12.45 | 24 | -36 | -44 |
| Insula | R | 4.86E-12 | 62 | 16.53 | 32 | 24 | 0 |
| Cerebellum Area 9 | L | 4.04E-14 | 82 | 13.71 | -10 | -50 | -46 |
| Cerebellum Area 10 | L |  |  | 13.41 | -22 | -36 | -42 |
| Thalamus | R | 7.99E-09 | 35 | 12.61 | 18 | -28 | 10 |
| **Salience Network** | | | | | | | |
| Dorsal Anterior Cingulate Cortex | R | <1E-15 | 9950 | 52.30 | 4 | 26 | 34 |
| Middle Frontal Gyrus | L |  |  | 48.21 | -30 | 46 | 28 |
| Middle Frontal Gyrus | R | <1E-15 | 2088 | 47.64 | 34 | 46 | 32 |
| Middle Frontal Gyrus | R |  |  | 23.38 | 42 | 44 | 14 |
| Insula | R | <1E-15 | 1433 | 45.89 | 36 | 18 | 8 |
| Insula | R |  |  | 42.09 | 40 | 14 | 2 |
| Insula | L | <1E-15 | 1934 | 45.40 | -38 | 14 | 6 |
| Insula | L |  |  | 45.34 | -32 | 18 | 10 |
| Supramarginal Gyrus | R | <1E-15 | 1342 | 39.77 | 62 | -36 | 38 |
| Supramarginal Gyrus | L | <1E-15 | 1074 | 38.46 | -60 | -40 | 34 |
| Inferior Parietal Gyrus | L |  |  | 12.85 | -52 | -44 | 56 |
| Cerebellum Crus I | R | <1E-15 | 465 | 27.84 | 46 | -54 | -30 |
| Cerebellum Crus I | R |  |  | 23.08 | 38 | -48 | -34 |
| Cerebellum Area 6 | L | <1E-15 | 183 | 25.69 | -34 | -52 | -32 |
| Cerebellum Crus I | L |  |  | 23.71 | -46 | -54 | -34 |
| Putamen | R | 1.41E-12 | 67 | 18.54 | 22 | 4 | 6 |
| Putamen | L | <1E-15 | 383 | 21.60 | -16 | 14 | -4 |
| Cerebellum Area 6 | L | 2.58E-14 | 84 | 21.90 | -40 | -52 | -26 |
| Middle Temporal Gyrus | L | <1E-15 | 210 | 20.10 | -54 | -64 | 0 |
| Middle Temporal Gyrus | R | <1E-15 | 184 | 19.77 | 54 | -56 | 0 |
| Pallidum | R | 5.78E-10 | 44 | 17.70 | 20 | 6 | 0 |
| Putamen | R |  |  | 13.03 | 14 | 12 | -8 |
| Precentral Gyrus | L | <1E-15 | 468 | 16.45 | -42 | -2 | 48 |
| Precentral Gyrus | L |  |  | 7.95 | -32 | 0 | 62 |
| Cerebellum Area 8 | L | 1.09E-08 | 34 | 15.34 | -32 | -44 | -44 |
| Thalamus | L | <1E-15 | 205 | 14.17 | -12 | -4 | 8 |
| Thalamus | R |  |  | 13.71 | 12 | -4 | 8 |
| Thalamus | L |  |  | 13.30 | -6 | -18 | 8 |
| Inferior Frontal Gyrus | R | 2.96E-12 | 64 | 11.41 | 24 | 32 | -16 |
| Insula | R |  |  | 10.85 | 26 | 24 | -12 |
| Precuneus | R | 1.15E-06 | 20 | 10.47 | 8 | -54 | 62 |
| **Sensorimotor Network** | | | | | | | |
| Superior Parietal Gyrus | L | <1E-15 | 30246 | 40.84 | -22 | -46 | 70 |
| Postcentral Gyrus | L |  |  | 40.25 | -36 | -40 | 64 |
| Postcentral Gyrus | R |  |  | 40.17 | 30 | -38 | 66 |
| Thalamus | R | 4.04E-08 | 31 | 12.96 | 12 | -18 | 6 |
| Thalamus | L | 2.86E-09 | 40 | 11.89 | -12 | -20 | 6 |
| **Visual Network** | | | | | | | |
| Calcarine | R | <1E-15 | 21772 | 52.49 | 10 | -88 | 4 |
| Calcarine | R |  |  | 51.86 | 6 | -80 | 2 |
| Fusiform Gyrus | R |  |  | 51.64 | 24 | -72 | -12 |
| Inferior Temporal Gyrus | R | 1.78E-06 | 20 | 13.33 | 54 | -52 | -22 |
| Coordinates are provided in MNI Space. Statistics are based on one sample t-test conducted on the individual maps extracted from the dual regression analyses. Age, Intracranial volume and site were added as covariates. | | | | | | | |

**6.1. Anatomical definition of the Atlas55+ RSNs’ subdivisions**

| **Supplementary Table S3: Anatomical Description of the RSN subdivisions** | |
| --- | --- |
| **Subdivision** | **Anatomical Description** |
| DMN Subdivision 1 (Green in Figure 4A) | Medial orbital prefrontal cortex |
|  | Precuneus/Posterior cingulate cortex |
|  | Bilateral superior prefrontal cortex |
|  | Bilateral hippocampi |
|  | Anterior middle temporal gyri |
|  | Bilateral angular gyri |
| DMN Subdivision 2 (Red in Figure 4A) | Anterior cingulate cortex |
|  | Medial orbital prefrontal cortex |
|  | Medial superior prefrontal cortex |
| DMN Subdivision 3 (Cyan in Figure 4A) | Left inferior frontal gyrus |
|  | Temporal poles |
|  | Supplementary motor area |
|  | Medial superior prefrontal cortex |
|  | Left middle temporal gyrus |
|  | Left precentral gyrus |
| DMN Subdivision 4 (Yellow in Figure 4A) | Left middle frontal gyrus |
|  | Left angular gyrus |
|  | Left medial superior prefrontal cortex |
|  | Left inferior/middle temporal gyrus |
| ECN Subdivision 1 (Purple in Figure 4B) | Right medial superior prefrontal cortex |
|  | Right middle/inferior frontal cortex |
|  | Right angular gyrus |
|  | Right middle/inferior temporal gyrus |
| ECN Subdivision 2 (Blue in Figure 4B) | Left medial superior prefrontal cortex |
|  | Left middle/inferior frontal cortex |
|  | Left inferior/superior parietal lobule |
|  | Left posterior inferior temporal gyrus |
| ECN Subdivision 3 (Green in Figure 4B) | Dorsal precuneus |
|  | Bilateral superior prefrontal cortex |
|  | Bilateral fusiform gyri |
|  | Bilateral parahippocampal gyri |
| ECN Subdivision 4 (Red in Figure 4B) | Bilateral inferior/superior parietal gyri |
| SAL Subdivision (Red in Figure 4C) | Bilateral supramarginal gyri |
|  | Bilateral inferior frontal cortex |
|  | Bilateral posterior middle temporal gyri |
|  | Dorsal anterior / middle cingulate cortex |
|  | Bilateral posterior insulas |
| SMN Subdivision 1 (Purple in Figure 4D) | Bilateral pre-/post-central gyri |
| SMN Subdivision 2 (Blue in Figure 4D) | Paracentral lobule |
|  | Supplementary motor area |
|  | Precuneus |
| SMN Subdivision 3 (Yellow in Figure 4D) | Left pre-/post-central gyri |
|  | Left supplementary motor area |
|  | Right precentral gyrus |
| SMN Subdivision 4 (Red in Figure 4D) | Bilateral superior temporal gyri |
| VIS Subdivision 1 (Blue in Figure 4E) | Bilateral lingual gyri |
|  | Bilateral cuneus |
|  | Bilateral calcarine |
| VIS Subdivision 2 (Red in Figure 4E) | Bilateral inferior/ Middle occipital cortex |
|  | Bilateral posterior calcarine |
| Anatomical description based on the Automated Anatomical Labelling (AAL) Atlas. | |

**6.3. Reliability of the major RSNs from Atlas55+**

In order to ensure the reproducibility of the Atlas55+ RSNs, we conducted several analyses. First, in order to endure that the spatial definition of the RSNs included in Atlas55+ was not influenced by our choice of approach (using the matlab function consensus_similarity.m), we constructed another version by assigning each brain voxel to a specific network only if at least 2 out of the 3 cohorts also assigned this voxel to the same network. The spatial correlation between the two versions of the Atlas55+ was very high with r-value above 0.9.

7. **Effect of age on the Atlas55+ RSNs**

| The effect of age on the spatial distribution of each RSN in the atlas (extracted from the dual regression) across all individuals (total n=563) was tested by conducting an analysis of covariance (ANCOVA), adding covariates of no interest: head motion, TIV, and site. Significant clusters are reported at a p<0.05 with FWE correction at the voxel level (T>5.3, cluster size>20 voxels, Supplementary Table S4).  **Supplementary Table S4: Clusters with significant association with age** | | | | | | | |
| --- | --- | --- | --- | --- | --- | --- | --- |
| **Regions** | **Hemisphere** | **P**  **(FWE-corr)** | **N of voxels** | **T** | **x** | **y** | **z** |
| **Negative Association with Age** | | | | | | | |
| **Default Mode Network** |  |  |  |  |  |  |  |
| Middle Cingulate Cortex | R | <1E-15 | 313 | 8.57 | 4 | -20 | 34 |
| Medial Orbitofrontal gyrus | R | <1E-15 | 169 | 7.41 | 2 | 52 | -12 |
| Medial Orbitofrontal gyrus | R |  |  | 5.78 | 4 | 40 | -10 |
| Superior Frontal gyrus | R | <1E-15 | 201 | 7.16 | 16 | 62 | 18 |
| Superior Medial Frontal gyrus | R |  |  | 6.99 | 10 | 58 | 36 |
| Anterior Cingulate Cortex | R/L | <1E-15 | 179 | 6.92 | 0 | 48 | 18 |
| Precuneus | R | <1E-15 | 290 | 6.54 | 8 | -66 | 30 |
| Posterior Cingulate Cortex |  |  |  | 6.35 | -10 | -46 | 32 |
| Middle Temporal gyrus | L | 1.59E-13 | 76 | 6.34 | -38 | -52 | 22 |
| pars orbitalis Inferior frontal gyrus | L | 3.72E-12 | 63 | 6.25 | -48 | 38 | -14 |
| Superior Frontal gyrus | L | 1.14E-06 | 20 | 5.68 | -18 | 58 | 26 |
| Middle Temporal gyrus | R | 1.96E-07 | 25 | 5.59 | 60 | -52 | 18 |
| **Executive Control Network** |  |  |  |  |  |  |  |
| Inferior Parietal Gyrus | L | <1E-15 | 250 | 8.67 | -38 | -46 | 38 |
| Inferior Parietal Gyrus | L |  |  | 5.90 | -48 | -38 | 38 |
| Middle Occipital Gyrus | L | <1E-15 | 191 | 7.71 | -26 | -64 | 30 |
| Middle Occipital Gyrus | L |  |  | 7.16 | -28 | -72 | 36 |
| Angular Gyrus | R | <1E-15 | 427 | 7.25 | 36 | -66 | 40 |
| Inferior Parietal Gyrus | R |  |  | 6.78 | 40 | -54 | 50 |
| Middle Frontal Gyrus | R | <1E-15 | 170 | 7.00 | 30 | 18 | 56 |
| pars opercularis inferior frontal gyrus | L | <1E-15 | 183 | 6.98 | -46 | 12 | 30 |
| pars opercularis inferior frontal gyrus | L |  |  | 6.14 | -36 | 8 | 26 |
| Superior Frontal gyrus | R | <1E-15 | 196 | 6.96 | 24 | 70 | 4 |
| Middle Frontal Gyrus | R |  |  | 6.38 | 36 | 64 | 4 |
| Middle Frontal Gyrus | R | 4.19E-13 | 72 | 6.27 | 38 | 34 | 22 |
| pars triangularis inferior frontal gyrus | R |  |  | 5.54 | 48 | 34 | 28 |
| pars opercularis inferior frontal gyrus | R | 1.81E-09 | 40 | 6.00 | 50 | 16 | 30 |
| Inferior Parietal Gyrus | R | 2.02E-08 | 32 | 5.93 | 50 | -40 | 50 |
| **Salience Network** |  |  |  |  |  |  |  |
| Dorsal Anterior Cingulate Cortex | L | <1E-15 | 953 | 9.82 | -2 | 28 | 26 |
| Dorsal Anterior Cingulate Cortex | R/L |  |  | 9.13 | 0 | 24 | 36 |
| Dorsal Anterior Cingulate Cortex | R |  |  | 7.79 | 8 | 26 | 30 |
| Anterior Insula | L | <1E-15 | 432 | 8.82 | -38 | 18 | 4 |
| Anterior Insula | L |  |  | 8.16 | -40 | 10 | -2 |
| Anterior Insula | R | <1E-15 | 305 | 8.42 | 38 | 10 | 2 |
| Anterior Insula | R |  |  | 6.65 | 34 | 22 | 8 |
| Middle Frontal Gyrus | L | <1E-15 | 300 | 7.85 | -34 | 54 | 16 |
| Superior Frontal gyrus | L | <1E-15 | 198 | 7.56 | -18 | 8 | 70 |
| Superior Frontal gyrus | L |  |  | 7.27 | -18 | 14 | 60 |
| Superior Frontal gyrus | R | <1E-15 | 150 | 6.98 | 22 | 10 | 66 |
| Supplementary Motor Area | R |  |  | 6.34 | 10 | 14 | 68 |
| Middle Cingulate Cortex | R/L | 1.29E-13 | 77 | 6.95 | 0 | -6 | 40 |
| Middle Frontal Gyrus | R | <1E-15 | 256 | 6.89 | 28 | 50 | 22 |
| Middle Frontal Gyrus | R |  |  | 6.48 | 32 | 58 | 24 |
| Middle Cingulate Cortex | L | 5.78E-10 | 44 | 6.87 | -8 | -20 | 42 |
| Supramarginal gyrus | L | 5.61E-07 | 22 | 6.17 | -56 | -42 | 34 |
| Supramarginal gyrus | R | 2.96E-12 | 64 | 5.99 | 66 | -38 | 36 |
| Inferior Parietal Gyrus | R |  |  | 5.94 | 56 | -36 | 48 |
| **Sensorimotor Network** |  |  |  |  |  |  |  |
| Superior Temporal gyrus | R | 7.14E-12 | 63 | 6.81 | 42 | -28 | 16 |
| Superior Temporal gyrus | L | <1E-15 | 167 | 6.78 | -54 | -24 | 18 |
| Superior Temporal gyrus | L |  |  | 6.15 | -52 | -32 | 20 |
| Postcentral gyrus | L | 5.2E-11 | 55 | 5.94 | -44 | -14 | 48 |
| Postcentral gyrus | L |  |  | 5.91 | -44 | -14 | 38 |
| Postcentral gyrus | R | 1.06E-06 | 21 | 5.84 | 38 | -16 | 36 |
| **Visual Network** |  |  |  |  |  |  |  |
| Lingual gyrus | R/L | 1.29E-12 | 72 | 6.76 | 0 | -76 | -6 |
| Calcarine gyrus | L |  |  | 5.82 | -4 | -92 | -8 |
| Lingual gyrus | R | 2.08E-08 | 34 | 6.32 | 10 | -92 | 26 |
| Calcarine gyrus | L | 9.36E-08 | 29 | 6.14 | -4 | -100 | 2 |
| Middle Occipital gyrus | L |  |  | 5.69 | -12 | -102 | 6 |
| **Positive Association with Age** | | | | | | | |
| **Default Mode Network** |  |  |  |  |  |  |  |
| Cerebellum - Crus I | R | 2.46E-10 | 47 | 6.31 | 30 | -76 | -30 |
| **Executive Control Network** |  |  |  |  |  |  |  |
| Cerebellum - Area 7b | L | 2.92E-11 | 55 | 6.46 | -32 | -64 | -44 |
| **Salience Network** |  |  |  |  |  |  |  |
| none |  |  |  |  |  |  |  |
| **Sensorimotor Network** |  |  |  |  |  |  |  |
| none |  |  |  |  |  |  |  |
| **Visual Network** |  |  |  |  |  |  |  |
| none |  |  |  |  |  |  |  |
| Coordinates are provided in MNI space. Only clusters with k>20 voxels and within the network of interest are shown. Height Threshold: T>5.3. | | | | | | | |
